# Spatial-filtering nanoscopy for 40-nm label-free Raman imaging

**DOI:** 10.64898/2026.09.03.749055

**Authors:** Jiye Li, Benma Lamu, Yuan Feng, Shengwen Wang, Haiyan Zhu, Zhiru Xu, Hulie Zeng

## Abstract

Super-resolution fluorescence microscopy overcomes the optical diffraction limit and has significantly advanced our understanding of biological complexity within the framework of fluorescence labelling^1^. Fluorescence labelling underpins this capability, enabling high photon budgets, superior signal contrast, and tuneable photophysical properties essential for diverse super-resolution modalities^2,3^. In contrast, label-free Raman imaging offers intrinsic chemical specificity^4^, supporting applications ranging from biomolecular fingerprinting to cell metabolic mapping^5–8^ and histopathological tissue characterization^9,10^. Despite its label-free advantage and chemical specificity, Raman imaging remains fundamentally limited in both spatial resolution and imaging contrast due to inherently low signal throughput and weak intrinsic Raman contrast^11–13^. Here, we introduce spatial-filtering nanoscopy (SFN), a physics-driven super-resolution strategy that achieves resolution enhancement through targeted signal purification instead of signal amplification, offering a conceptually distinct pathway beyond the diffraction limit. SFN synergistically integrates a sub-millimetre microsphere lens (SMML) with a standard confocal Raman microscope, harnessing two complementary physical effects: (i) the photonic redistribution effect (PRE), which narrows the lateral excitation profile, and (ii) the three-dimensional spatial filtering (3D-SFE), which effectively suppresses both lateral and axial background. We demonstrate SFN-enabled super-resolution Raman imaging of silicon nanostructures, intact cells, and tissue sections, achieving an effective lateral resolution of approximately 40 nm, and chemically resolving subcellular features including organelles and pseudopodia without exogenous labels. These findings establish SFN as a broadly generalizable hardware-based super-resolution strategy, readily deployable on standard confocal platforms and extensible across diverse optical imaging modalities.

## Principle and performance characterization of SFN system

The SFN system was constructed by integrating a commercially available K9 SMML with a diameter of 500 μm into a conventional confocal Raman microscope (CRM). The SMML was placed directly on the sample surface and immersed in water to optical match with a 63× water-immersion objective with a numerical aperture (NA) of 1.0). Under this configuration, Raman imaging was performed using 532 nm excitation (Fig. 1a). Analogous to microsphere-assisted microscopy (MAM)^14^, the SMML generates a magnified virtual image on the sample side. The theoretical magnification SMML could be given by (Supplementary Note 1):

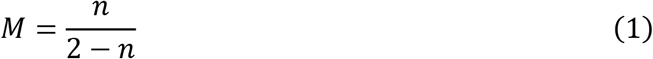

**Fig. 1.**
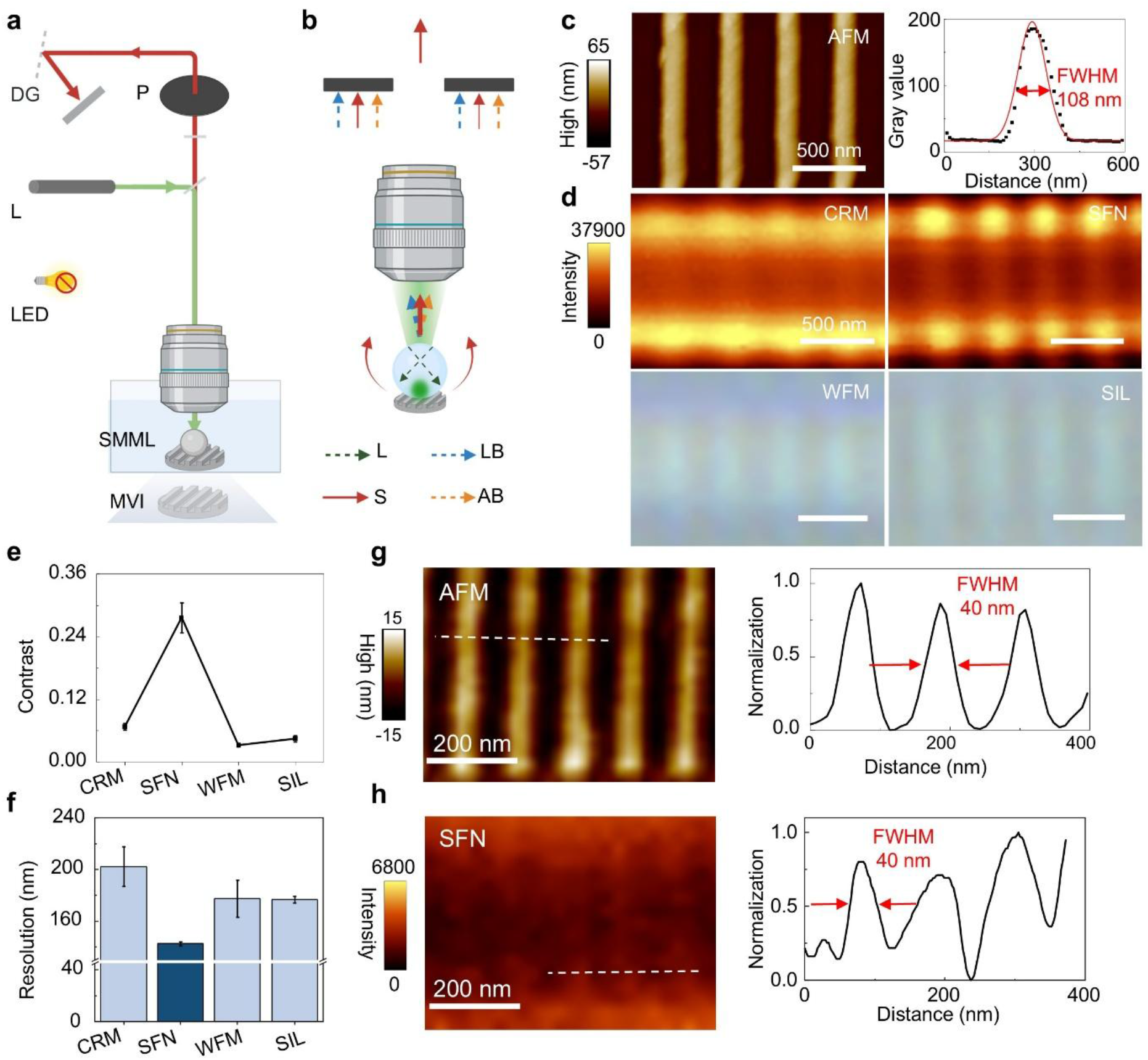
Principle and characterization of the SFN system. **a**, Schematic illustration of the SFN configuration. MVI, magnified virtual image; L, 532 nm laser; P, pinhole; DG, diffraction grating; LED, white-light source for wide-field imaging. **b**, Conceptual mechanism of SFN resolution improvement. SFN achieves improved spatial resolution through concurrent suppression of lateral background (LB) and axial background (AB), while preserving the useful signal (S). Line widths in the schematic are proportional to relative intensity. **c**, AFM image of the silicon nanostructure (FWHM = 108 ± 2 nm) used for resolution assessment. **d**, Images of the nanostructure acquired using CRM (upper left), SFN (upper right), WFM (lower left) and SIL (lower right). **e**, Michelson contrast measured for each imaging mode in **d**. **f**, Apparent linewidths extracted from Gaussian fitting of the images in **d**. **g**, AFM image of another silicon nanostructure (FWHM = 40 ± 1 nm) used for evaluation of imaging quality. **h**, SFN Raman image of the approximately 40 nm nanostructure. Acquisition parameters of all Raman imaging: 30 nm step size, 520 cm⁻¹ Si-Si mode, 0.5 s integration time and 20 mW laser power. All linewidths were determined by Gaussian fitting of line profiles and FWHM was used as the resolution metric. Data are presented as mean ± s.d. over the field of view (FOV).

where *M* is magnification and *n* = *n*_SMML_/*n*_A_ is the relative refractive index of the SMML, with *n*_SMML_ and *n*_A_ denoting the refractive indices of the SMML and the ambient medium, respectively. The results indicate that an effective magnification of approximately 1.32× could be obtained relative to that of Raman imaging acquired through a bare 63×/1.0 NA objective. This magnification is independent of the physical size of the SMML and depends solely on the refractive indices of the SMML and the ambient medium (Supplementary Fig. 1).

The SFN system leverages two synergistic physical mechanisms: photonic redistribution effect (PRE) and three-dimensional spatial filtering effect (3D-SFE). PRE compresses the lateral excitation profile and reshapes the emission pattern, whereas 3D-SFE suppresses both lateral and axial background contributions (Fig. 1b). A conventional CRM yields a diffraction-limited lateral resolution of approximately 325 nm. When coupled with the SMML, PRE alone improves the nominal lateral resolution from approximately 325 nm to 214 nm; subsequently 3D-SFE introduces additional lateral filtering, further reducing the nominal resolution to approximately 94 nm. The combination of PRE and 3D-SFE yields a nominal lateral resolution of approximately 86 nm. Moreover, 3D-SFE also strongly suppresses the axial background, a mechanism previously demonstrated to enhance lateral resolvability in confocal microscopy^15,16^, thereby pushing the effective lateral resolution down to approximately 40 nm. Concurrently, the dual suppression of PRE and 3D-SFE significantly enhances image contrast, collectively improving overall imaging fidelity.

To quantitatively evaluate the imaging performance of the SFN system, we imaged a silicon nanostructure with a full-width at half-maximum (FWHM) of 108 ± 2 nm, as measured by atomic force microscopy (AFM) and verified by Gaussian fitting (Fig. 1c). Reference images of the same nanostructure were acquired using conventional CRM, bright-field wide-field microscopy (WFM) and SMML-coupled wide-field microscopy (SIL)^17^. The contrast of each imaging modality was quantified using the Michelson contrast (Fig. 1d). Specifically, the images acquired through SFN exhibited a Michelson contrast of 0.28 ± 0.006, compared with that of 0.03 ± 0.001 for WFM, 0.04 ± 0.006 for SIL and 0.07 ± 0.006 for CRM (Fig. 1e). Thus, SFN delivers approximately a fourfold contrast enhancement over conventional CRM, attributable to the combined lateral and axial background suppression provided by PRE and 3D-SFE. Furthermore, Gaussian fitting of the Raman image acquired by SFN yielded an FWHM of 143 ± 2 nm for the nanostructure. In comparison, the apparent linewidths of the same nanostructure were measured to be 202 ± 15 nm, 177 ± 16 nm and 177 ± 3 nm via CRM, WFM and SIL, respectively (Fig. 1f). This result demonstrates that the substantially improved spatial resolution achieved by SFN originates from the synergistic effects of PRE and 3D-SFE. Notably, the measured linewidths of the nanostructure via CRM, WFM, and SIL were all narrower than their respective theoretical diffraction limits, likely due to the exceptionally sharp and high-contrast morphology of the nanostructure. Additionally, the SFN Raman image of the nanostructure gave an FWHM of 143 ± 2 nm, broader than the AFM reference, indicating that the effective resolution is constrained by background signals that blur the nanostructure edges.

To further validate the resolving capability of the SFN system, we used the other silicon nanostructure with an AFM-measured FWHM of 40 ± 1 nm (Fig. 1g). Raman imaging of the same feature via SFN gave a minimum FWHM of ∼40 nm from Gaussian fitting (Fig. 1h), confirming that the effective lateral resolution of the SFN system is at least 40 nm.

## Qualitative and quantitative analysis of signal purification of SFN

To qualitatively and quantitatively characterize the synergistic physical effects underlying SFN, we primarily developed a comprehensive theoretical framework that decomposes the total signal enhancement ratio (*E*_total_) into three independent physical components: laser coupling efficiency (*η*_c_), focusing gain (*G*_f_), and signal collection efficiency (*E*_col_). We first verified that both the SFN and SIL systems operate within the linear response regime (Extended Data Fig. 1 and Supplementary Note 2), consequently, the decomposition of *E*_total_ imparted by the SMML upon coupling with the microscope into above three independent components expressed as (Supplementary Note 3):

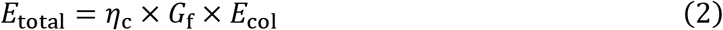

To qualitatively and quantitatively clarify the donation of PRE in SFN system, we isolated the SIL system without confocal pinhole (dashed box, Fig. 2a). The *E*_total_ was measured on a polished silicon wafer under 532 nm illumination. At this stage, the axial contribution was negligible, so that the system could be considered as a two-dimensional (2D) model. By normalizing the gray value of the first-order Airy bright ring obtained under the SIL configuration that measured under at the incident excitation power of 0.10 mW referring that of the WFM system without SMML coupling, we observed an approximately 1.5-fold increase in the effective excitation intensity upon SMML integration, indicating that the *E*_total_ of the SIL configuration for SMML coupling is equivalent to a 1.5-fold enhancement (Fig. 2b).

**Fig. 2.**
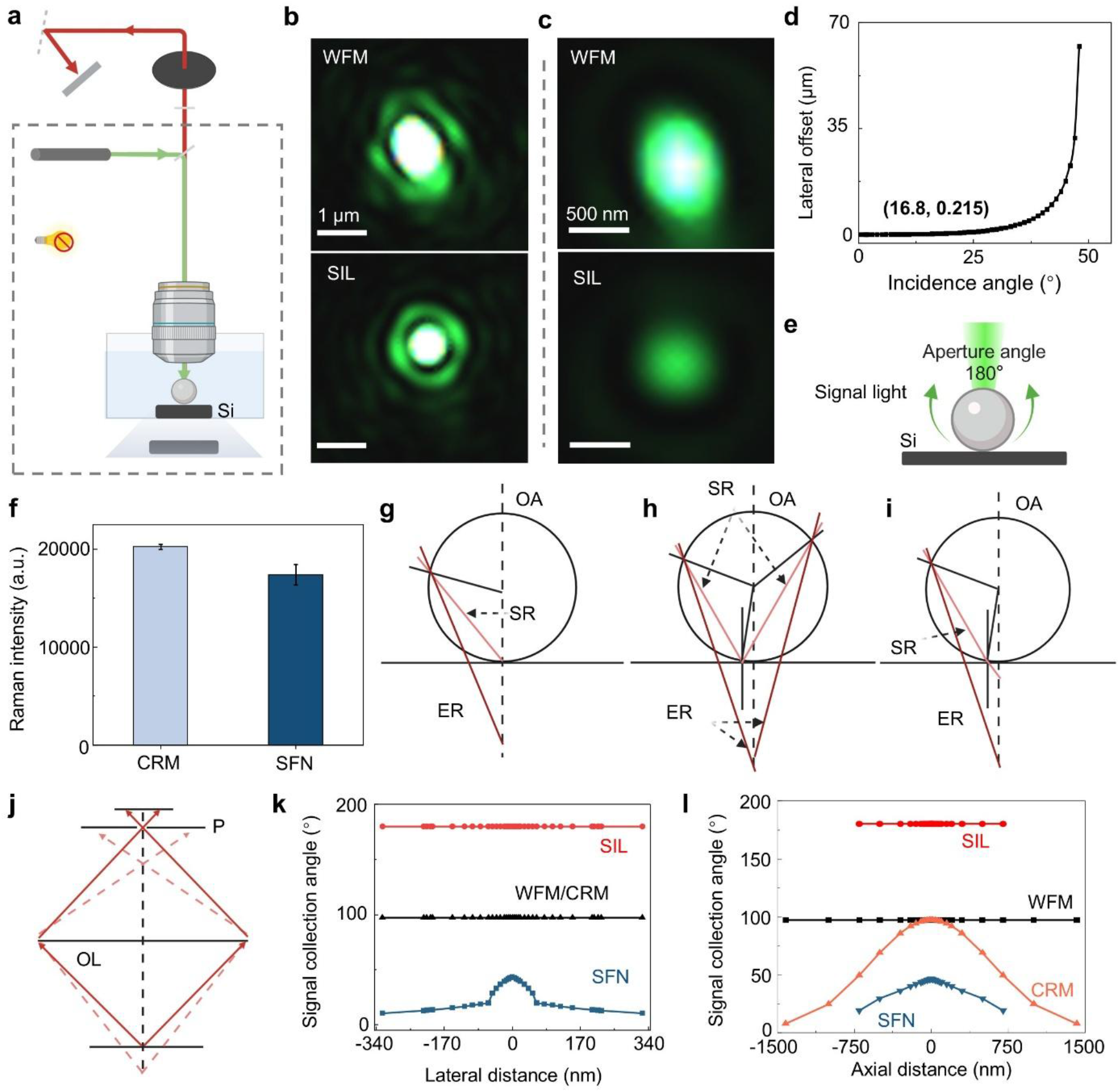
Qualitative and quantitative analysis of signal purification of SFN. **a**, Schematic of optical configuration of SFN. The dashed square is the optical configuration used for PRE analysis. **b**, Experimental Airy disk images acquired under WFM and SIL systems using 0.1 mW 532 nm laser illumination. **c**, Airy disk images acquired under dual-source illumination (0.1 mW, 532 nm laser + 0.1% white-light). **d**, Lateral offset of the focused spot within the SMML as a function of incident laser angle. **e**, Schematic of the effective aperture angle in SIL geometry. **f**, Quantitative comparison of Raman signal intensities from a polished silicon wafer, measured at the Si-Si stretching mode (520 cm⁻¹), under identical acquisition parameters (20 mW laser power, 0.5 s integration time, 10 accumulations). Data are presented as mean ± s.d. (N = 5). **g-i**, Equivalent geometric-optics models underpinning SFN performance: (**g**) focal-point model; (**h**) lateral-offset model; (**i**) axial-offset model. SR, signal ray; ER, equivalent ray; OA, optical axis. **j**, Schematic illustrating the axial filtering effect of conventional CRM. **k**, **l**, Theoretically derived lateral (**k**) and axial (**l**) angular distributions of collected signals for WFM, CRM, SIL, and SFN configurations.

To isolate the *G*_f_ component, we characterized the Airy disk morphology generated in the SIL configuration. A dual-source illumination strategy was implemented to prevent saturation while enabling precise measurement of the central Airy spot shape and diameter (Fig. 2c and Supplementary Note 4). SMML coupling transformed the excitation focus from an elliptical profile to a near-circular one, reducing the major-to-minor axis ratio from 1.3 ± 0.1 to 1.1 ± 0.1 (Supplementary Fig. 2a). Concurrently, the major axis decreased from 644.0 ± 35.3 nm to 429.6 ± 9.2 nm (Supplementary Fig. 2b), indicating substantial mitigation of focal anisotropy and axial elongation. Translating this improvement into effective numerical aperture (NA_eff_), the SIL system achieved NA_eff_ = 1.51 ± 0.03, compared to 1.01 ± 0.05 for the bare WFM, corresponding to a *G*_f_ of 2.24×.

Having quantified *G*_f_, we next determined *η*_c_ and *E*_col_ using a geometric optics model applicable to both laser coupling and signal collection. Ray-tracing simulations revealed that approximately 35% of paraxial laser rays (approximately 33.6° out of a total 97.2°) are effectively focused into the SIL system (Fig. 2d and Supplementary Note 5). Additionally, the SIL system collects signal over a nearly 90° aperture half-angle, yielding an approximately 1.85× enhancement in *E*_col_ (Fig. 2e and Supplementary Note 6).

Multiplying the individual contributions according to Eq. (2), *η*_c_ = 0.35, *G*_f_ = 2.24, and *E*_col_ = 1.85, yields the theoretical *E*_total_ ≈ 1.5×, in excellent agreement with experimentally measured values. This consistency between theoretical deduction and experimental measured signal enhancement *E*_total_ also well proves the validity of PRE-based theoretical framework and Eq. (2). Therefore, we extend the analytical framework of PRE to SFN system since SFN share an identical excitation pathway with SIL system, and the PRE contributions (*η*_c_ and *G*_f_) remain unchanged. The only difference between SFN system with SIL system lies in the detection pathway, where SFN additionally incorporates the confocal pinhole and thus introduces 3D-SFE. To well characterize the 3D-SFE, we dissected it into lateral and axial components. Lateral filtering was isolated and investigated by acquiring Raman spectra from a polished silicon wafer under identical excitation conditions across SFN and conventional CRM configurations (Fig. 2a). The Raman signal intensity measured in SFN was substantially lower than that in CRM, directly evidencing the lateral filtering effect of SMML coupling. To further elucidate this mechanism, we developed a qualitative geometric optics model that maps the SFN system to the well-established CRM framework through equivalent axial positions and emission angles (Fig. 2g-i and Supplementary Note 7). The model reveals that the SMML redirects signal rays toward the axial direction, and that 3D-SFE arises as a consequence of SMML-pinhole coupling.

Building on this insight, we developed a point-spread function (PSF) model to quantitatively evaluate the 3D-SFE (Fig. 2j and Supplementary Note 8). Comparative analysis of lateral signal collection across platforms reveals distinct behaviors: (i) conventional WFM and CRM exhibit uniform lateral collection; (ii) the SIL system enhances signal collection efficiency while preserving lateral uniformity; and (iii) SFN introduces pronounced lateral filtering, which selectively suppresses lateral background while only moderately attenuating the focal signal (Fig. 2k). This trade-off, namely significant background suppression with modest signal loss, is fundamental to lateral filtering. Analogous analysis of axial signal collection confirms enhanced axial filtering in SFN versus CRM (Fig. 2l). We can see Eq. (2) is effective for the PRE model at 2D framework, more models and deduction are necessary to the 3D frameworks.

## Quantitative analysis of signal intensity of SFN

To enable quantitative cross-platform comparison of signal intensity, we constructed a unified theoretical model that incorporates both the excitation and collection characteristics of each imaging configuration. Specifically, the excitation intensity distributions of all configurations were normalized to that of the WFM reference, and configuration-specific collection angle distributions were incorporated. This yielded per-platform signal intensity distributions, which were then converted into relative signal intensities for direct cross-platform comparison (Supplementary Notes 9 and 10).

Under the 2D framework, we computed the *E*_total_ for each configuration using WFM as the reference (Extended Data Fig. 2a). We first investigated SIL system to validate the theoretical model of 2D framework, the *E*_total_ ≈ 1.5×was calculated and agreed well with our earlier experimental measurements, confirming the reliability of our modelling approach. We then extended the theoretical model of 2D framework to CRM and SFN, where a substantial discrepancy between theoretical and experimental results emerged. We attribute this discrepancy to the coupling between the signal angular distribution and the grating efficiency. Although the 1.0 NA objective collects photons over a wide angular range, high-angle rays are diffracted with lower efficiency. The SMML compresses the angular spread, thereby reducing grating loss and partially offsetting the predicted signal decrease.

We further probed the influence of SMML diameter and refractive index on signal intensity using the same 2D framework. Simulations accurately reproduce the experimentally observed trends for both parameters, namely a gradual decrease in signal intensity with increasing SMML diameter and refractive index (Supplementary Fig. 3 and 4). Deviations in several measurements, particularly for the 300-μm K9 and 500-μm fused silica SMMLs, likely arise from fabrication imperfections that reduce the actual optical coupling efficiency. Additionally, residual mismatches suggest that angle-dependent losses at the diffraction grating affect the detected signal, consistent with the grating-efficiency effect discussed above.

**Fig. 3.**
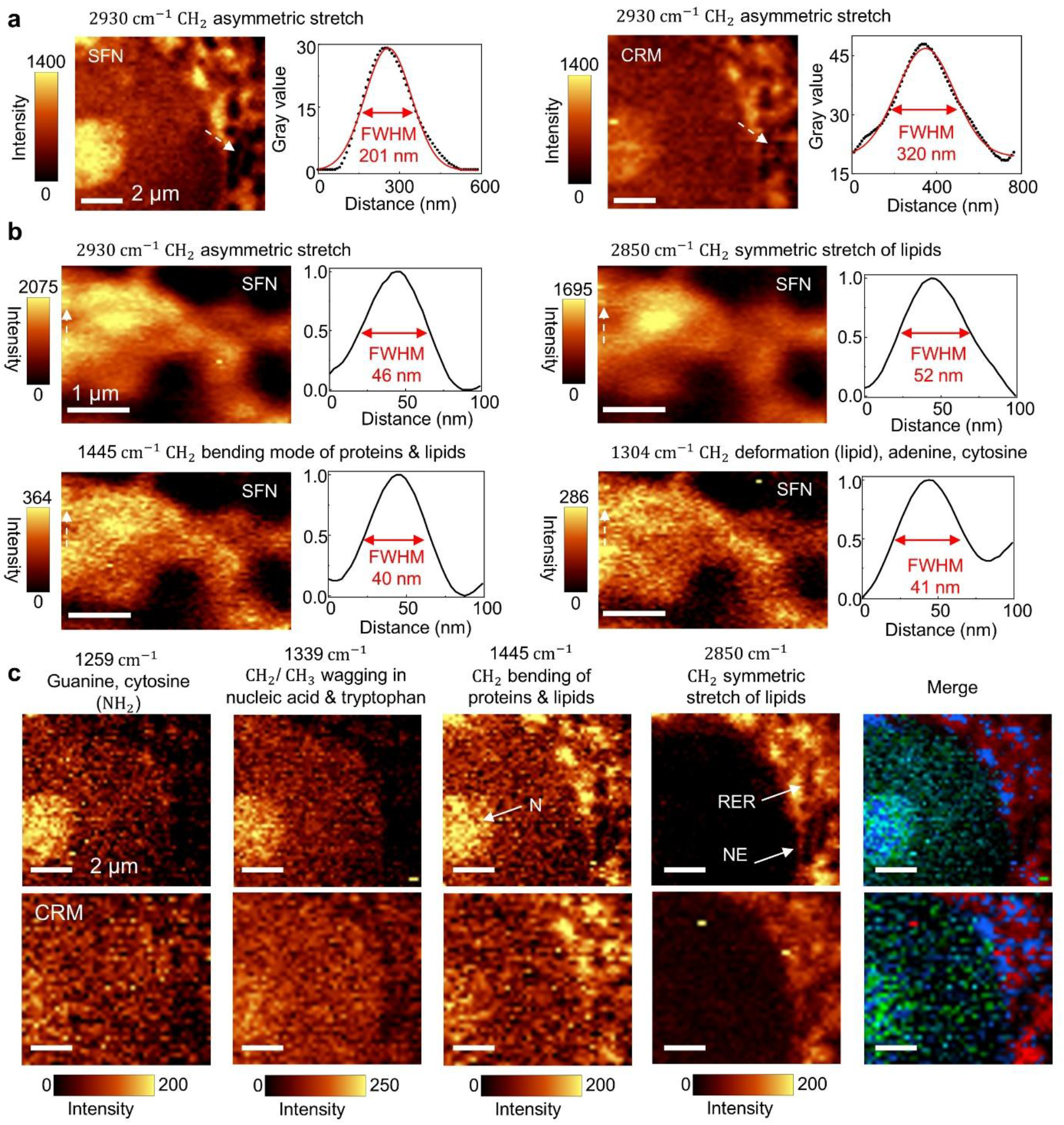
Super-resolution Raman imaging of A549 cells. **a**, Comparative Raman images of a representative cellular region acquired at 2930 cm⁻¹: conventional CRM (left) and SFN (right). Corresponding lateral intensity profiles (dashed lines) are overlaid to the right of each image. **b**, Multispectral SFN Raman images at four bands: 2930 cm⁻¹, 2850 cm⁻¹, 1445 cm⁻¹, and 1304 cm⁻¹, with corresponding lateral intensity profiles. **c**, Raman images of a nuclear region showing four channels and their pseudocolor merge: CRM (top row) and SFN (bottom row). Four channels correspond to 1259 cm⁻¹, 1339 cm⁻¹, 1445 cm⁻¹, and 2850 cm⁻¹. Acquisition parameters: 200 nm step size in **a** and **c**, 30 nm step size in **b**; 20 mW laser power and 1.0 s integration time. Annotations: N, nucleus; NE, nuclear envelope; RER, rough endoplasmic reticulum.

**Fig. 4.**
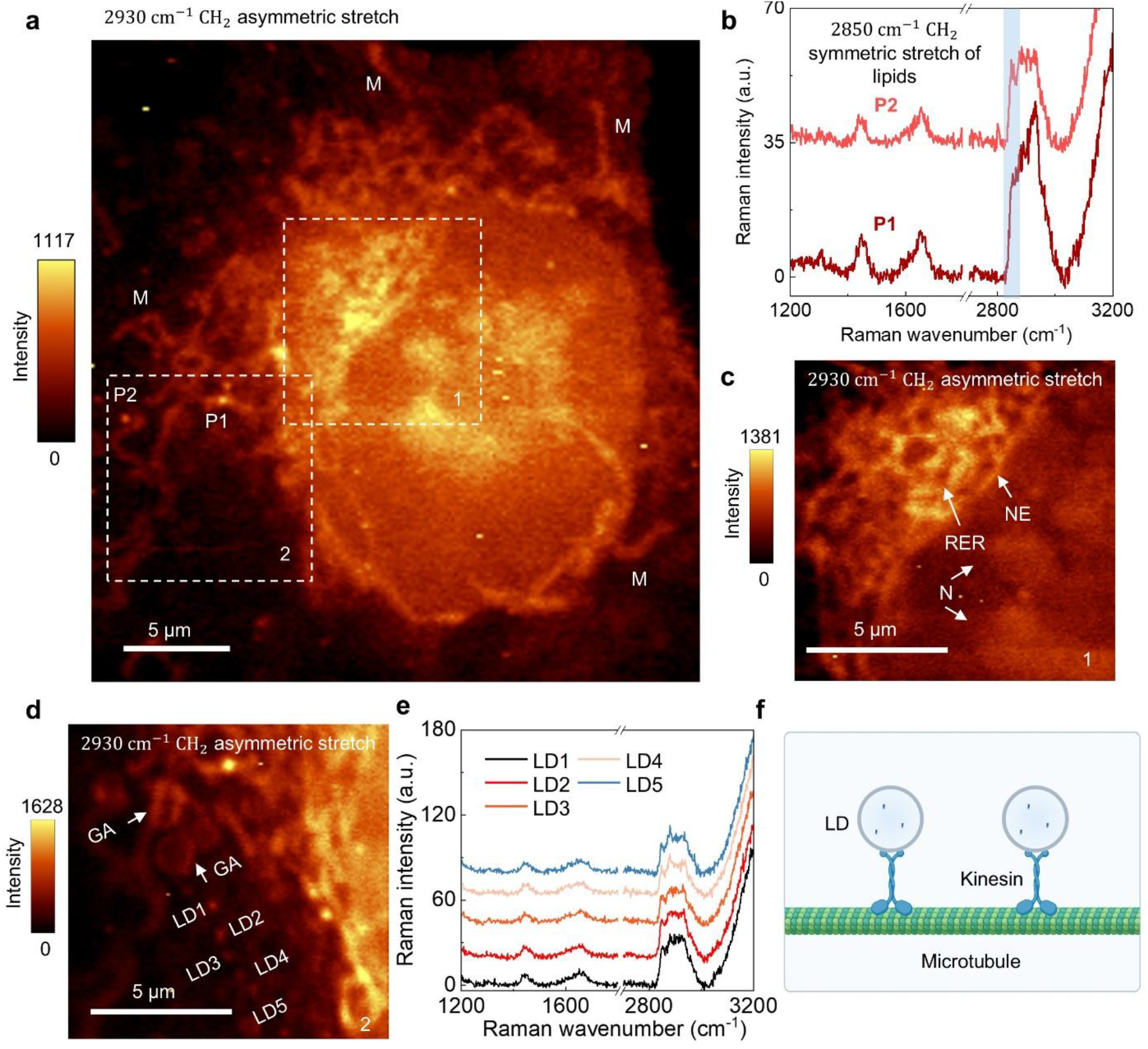
Nanoscale Raman imaging of subcellular organelles in HUVECs. **a**, Large field-of-view Raman image of HUVECs acquired at 2930 cm⁻¹, with a step size of 200 nm. Mitochondria (M) are identified based on their characteristic perinuclear distribution and morphology. **b**, Representative Raman spectra extracted from spatially distinct regions P1 and P2 in **a**, highlighting differential lipid signatures. P1 exhibits lower lipids signal intensity (2850 cm^-1^) consistent with an early endosomes or transport vesicles, whereas P2 shows elevated signal indicative of lipids enrichment. **c**, High-resolution Raman image of ROI-1 acquired at 2930 cm⁻¹ with a 100 nm step size. Structural features resolved include the reticular architecture of RER, N, and NE. **d**, High- resolution Raman image of region of interest 2 (ROI-2), acquired at 2930 cm⁻¹ with a 100 nm step size. The flattened cisternal morphology characteristic of the Golgi apparatus (GA) is clearly delineated. **e**, Raman spectra (2930 cm⁻¹ channel) extracted from five linearly aligned LDs identified within ROI-2, confirming their uniform lipid-rich composition. **f**, Schematic illustration of the spatial arrangement of the five LDs along a linear trajectory, consistent with directed transport along microtubules. Acquisition parameters: 20 mW laser power, 1.0 s integration time.

Finally, we extended the analysis to 3D systems by incorporating axial background contributions into the model. Using the relative signal intensity of the 2D WFM as the reference, we calculated the *E*_total_ for all configurations under the 3D framework (Extended Data Fig. 2b). In this 3D framework, the WFM signal was substantially increased compared with the 2D case due to the inclusion of out-of-focus background. Relative to the WFM under 3D framework, conventional CRM exhibited a modest reduction in signal intensity, attributable to the axial filtering effect of the confocal pinhole. The SIL configuration also showed a slight decrease in signal intensity, attributable to PRE-induced focal spot compression and excitation loss, with a combined factor of *η*_c_ × *G*_f_ ≈ 0.8. This reduction decreases the effective excitation volume and consequently suppresses the integrated axial signal. In contrast, SFN did not yield a significant signal enhancement over CRM. This observation indicates that SFN possesses pronounced axial filtering capability, effectively suppressing out-of-focus background while preserving the in-focus signal.

## Theoretical analysis of resolution and signal-to-background ratio of SFN

To rigorously assess the resolution limit of SFN, we performed theoretical calculations grounded in the PSF separation principle, according to which overall spatial resolution is jointly determined by excitation confinement and collection filtering (Supplementary Note 11). Specifically, PRE alone yields an excitation-limited lateral resolution of approximately 214 nm, while lateral filtering refines the collection-limited resolution to approximately 94 nm (Extended Data Fig. 3a). The convolution of these two contributions predicts an overall theoretical lateral resolution of approximately 86 nm (Extended Data Fig. 3b). Notably, the experimentally measured resolution reaches approximately 40 nm, significantly surpassing this prediction. We attribute this enhancement to the strong suppression of axial background by 3D-SFE, consistent with previous reports showing that reducing axial background intensity improves lateral resolvability^15,16^. Accordingly, we quantified the signal-to-background ratio (SBR) across all imaging configurations (Extended Data Fig. 3c and Supplementary Note 12). In the 2D model, SBR increases from 0.40 for both WFM and CRM to 0.73 for SIL (a 1.8-fold improvement) and further to 1.50 for SFN (a 3.8-fold improvement over WFM and CRM), with PRE and lateral filtering acting as the primary drivers, respectively. In the 3D model, SFN achieves an SBR of 0.75, representing an 8.4-fold enhancement relative to WFM, and substantially exceeding the gains achieved by SIL (4.5-fold) and CRM alone (1.5-fold). These results demonstrate that SFN possesses superior axial filtering capability, which suppresses background and substantially improves imaging contrast, thereby enabling further enhancement of lateral resolution.

## Effective numerical aperture of SFN

NA is a fundamental parameter governing both optical resolution and signal collection efficiency in microscopy. Based on our geometric-optics model, the effective aperture half-angle of the SFN system approaches 90° upon SMML integration. When the SMML is coupled with a high-NA objective, the NA_eff_ is predominantly determined by the refractive index of SMML, as described by Eq. (3):

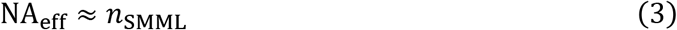

Where *n*_SMML_ denotes the absolute refractive index of the SMML material (Supplementary Note 13). Consequently, the SFN system functions analogously to an oil-immersion objective. The SMML serves partly as the high-index immersion medium and partly as an integral component of the objective (Supplementary Fig. 5). This analogy provides an intuitive interpretation of the observed behaviour.

## PRE-modulated signal intensity in SIL and SFN

Signal intensity is a critical performance metric for SFN, as it directly governs both the SBR and the signal-to-noise ratio (SNR). Among the factors influencing signal intensity, PRE serves as the key modulator of both excitation and signal collection. Laser coupling efficiency is the primary factor that limits both the excitation and the signal intensity under PRE regulation. Our theoretical analysis reveals that this coupling efficiency is most sensitively dependent on the refractive index of the ambient medium and the NA of the objective (Extended Data Fig. 4a, b), whereas the refractive index and diameter of SMML and excitation wavelength exert comparatively weaker influences (Extended Data Fig. 4c-e). These insights rationalize the experimental choice of a water-immersion objective and establish a generalizable design framework for optimizing SIL and SFN performance across diverse imaging configurations.

## Biocompatibility and photodamage mitigation of SFN

Conventional super-resolution microscopy techniques often rely on high excitation intensities and prolonged acquisition times, conditions that mechanistically induce photodamage and thus pose a significant challenge for high-resolution imaging of delicate biospecimens^18,19^. In contrast, the SFN system enables super-resolution Raman imaging with reduced excitation burden and efficient thermal diffusion, thereby minimizing photothermal damage to the specimen. Specifically, SMML integration reduces the effective excitation intensity at the focal plane to approximately 0.8 (*η*_c_ × *G*_f_ ≈ 0.8). This feature, together with the favourable thermal properties of K9 glass, helps mitigate photodamage and supports long-term cell imaging.

## Spectral performance of SFN

High spectral fidelity is essential for resolving subtle molecular distributions in high-resolution Raman imaging of biological specimens, particularly given the intrinsically weak Raman cross-sections of biomolecules and their susceptibility to background interference^20,21^. To systematically evaluate how SMML material choice affects spectral fidelity, we characterized the intrinsic Raman responses of candidate microspheres using a silicon wafer as a standardized reference (Supplementary Note 14). The K9 SMML employed in SFN exhibits negligible intrinsic Raman background across the measured spectral range and preserves quantitative linearity between incident excitation power and detected Raman signal intensity (Extended Data Fig. 1), confirming the suitability of SFN for quantitative Raman imaging. In contrast, background and noise intensity displayed a pronounced nonlinear dependence on excitation power (Extended Data Fig. 5b,c), consistent with laser-induced thermal radiation^22^. Critically, relative to conventional CRM, SFN suppresses spectral background while maintaining comparable noise levels. This significantly enhances spectral contrast and enables reliable detection of low-intensity Raman features that would otherwise be obscured (Extended Data Fig. 5c).

## Super-resolution imaging of biospecimens by SFN

Label-free Raman imaging of biospecimens at high resolution has remained a longstanding challenge, primarily because biomolecules possess intrinsically weak Raman scattering cross-sections, typically orders of magnitude lower than those of inorganic materials such as crystalline silicon^23,24^. This fundamental signal paucity imposes a severe trade-off: achieving higher spatial resolution demands finer sampling and longer integration times, yet these conditions do not circumvent the diffraction limit; they merely accumulate more of the same low-signal, high-background data. To rigorously evaluate whether SFN can break this deadlock, we performed comparative Raman imaging of intact A549 human lung adenocarcinoma cells using both the SFN system and conventional CRM under identical imaging conditions.

At the CH₂ asymmetric stretching band (2930 cm⁻¹) ^25^, SFN achieved a lateral resolution of 200 nm with a 200-nm step size, matching the confocal sampling interval (Fig. 3a). In contrast, CRM acquired under identical parameters yielded only 320 nm resolution over the same cellular region, consistent with the theoretical diffraction limit (approximately 325 nm). These results demonstrate that SFN enables true super-resolution Raman imaging in intact biological cells, whereas CRM remains fundamentally constrained by the optical diffraction barrier.

To quantitatively assess the sub-diffraction resolving power of SFN, we acquired Raman imaging of rough endoplasmic reticulum (RER) in A549 cells using a 30-nm step size. Four characteristic CH₂ vibrational modes were independently resolved: 2930 cm⁻¹ (asymmetric stretch), 2850 cm⁻¹ (symmetric stretch of lipids), 1445 cm⁻¹ (bending), and 1304 cm⁻¹ (deformation)^25^. SFN achieved a minimum lateral resolution of approximately 40 nm across these bands (Fig. 3b). Notably, resolution exhibited a modest dependence on vibrational mode intensity: higher-intensity bands (2930 and 2850 cm⁻¹) yielded slightly coarser resolutions (46 nm and 52 nm, respectively), whereas weaker bands (1445 and 1304 cm⁻¹) achieved superior resolution (40 nm and 41 nm). This inverse correlation arises because vibrational modes with higher intrinsic signal intensities are accompanied by higher background levels, reducing the effective SBR and thereby limiting resolvability. Moreover, the background suppression capability of SFN substantially enhances the contrast of weak Raman signals, enabling reliable subcellular structural mapping that is inaccessible to conventional CRM (Fig. 3c). This contrast enhancement stems from two independent contributions: 3D-SFE suppresses out-of-focus background, while PRE compresses the excitation spot and improves the utilization efficiency of incident laser power, thereby boosting the detectable Raman signal from cellular specimens. In the inherently weak-signal regime of cellular Raman imaging, both mechanisms are essential for achieving the observed resolution and contrast gains.

We next examined whether the resolution gains observed in cultured cells translate directly to tissue specimens. Cryosections of mouse lung tissue imaged via SFN exhibited analogous resolution improvements (Extended Data Fig. 6). However, the contrast improvement relative to CRM was markedly attenuated in tissues compared with cultured cells. This discrepancy stems from the greater thickness and higher macromolecular density of tissue specimens, which enhance the effective excitation volume and Raman signal intensity in CRM. Consequently, the contrast gain conferred by SFN is inherently reduced in tissues, whereas the weaker intrinsic signals in cultured cells allow the background-suppression and signal-enhancement mechanisms of SFN to manifest more robustly.

To corroborate this interpretation, we quantitatively compared Raman signal performance between the two specimen types using spectra extracted from their respective Raman images (Extended Data Fig. 7). In cultured cells, SFN consistently delivered higher Raman signal intensities than CRM, coupled with significant suppression of spectral background and noise, improving both SNR and SBR. In tissue sections, by contrast, SFN did not yield net signal enhancement over CRM. Although 3D-SFE continues to suppress spatial background, the overall signal intensity remains unchanged because the dense tissue architecture elevates baseline spectral background and noise, partially offsetting the filtering benefit. Collectively, these results demonstrate that SFN is particularly advantageous for cell imaging, where it simultaneously enhances both SNR and SBR.

## Super-resolution Raman imaging of subcellular organelles by SFN

To further evaluate the feasibility and analytical utility of SFN for nanoscale subcellular Raman imaging, we performed high-resolution Raman imaging of organelles in human umbilical vein endothelial cells (HUVECs). Initial large field-of-view Raman imaging at a 200-nm step size resolved a mitochondrion distributed throughout the cytoplasm and encircling the nucleus, identified by its morphology and characteristic spatial distribution at 2930 cm⁻¹ (Fig. 4a). Two distinct subcellular regions, designated P1 and P2, exhibited differential Raman intensities at 2850 cm⁻¹, assigned to lipid acyl chains. This contrast reflects regional heterogeneity in lipid content (Fig. 4b): P2 displayed significantly higher signal intensity than P1, consistent with the biochemical signature of lipid droplets (LDs), whereas P1 corresponded to a less lipid-enriched vesicular compartment, likely early endosomes or transport vesicles.

To achieve finer structural discrimination, we performed targeted high-resolution Raman imaging on two regions of interest, ROI-1 and ROI-2, with a reduced step size of 100 nm. In ROI-1, the reticular architecture of the rough endoplasmic reticulum (RER), along with the nucleus and nuclear envelope, was unambiguously resolved (Fig. 4c). In ROI-2, the stacked, flattened cisternal morphology characteristic of the Golgi apparatus was clearly visualized (Fig. 4d). The increased spatial sampling density markedly improved structural definition and enabled precise delineation of organelle boundaries. Within ROI-2, five linearly aligned LDs were identified; their spatial distribution profiles at 2850 cm⁻¹ confirmed their lipid-rich composition (Fig. 4e). Notably, the colocalization of these LDs along a linear trajectory is strongly indicative of active, microtubule-dependent intracellular transport (Fig. 4f)^26^. This observation demonstrates the capacity of SFN to resolve spatially organized, biochemically specific features that directly report on functional intracellular processes.

## Super-resolution Raman imaging of pseudopodial lipid metabolism by SFN

Pseudopodia are transient, actin-driven cytoplasmic protrusions that dynamically coordinate with LDs and the endoplasmic reticulum during cellular adaptation^27^. However, their nanoscale metabolic heterogeneity, particularly in lipid composition, spatial distribution, and trafficking dynamics, has remained inaccessible to conventional Raman microscopy techniques^28^. This is largely because pseudopodia are substantially thinner than the cell body, yielding intrinsically weaker Raman signals that fall below the detection sensitivity of diffraction-limited CRM. To overcome this limitation, we performed nanoscale subcellular Raman imaging of lipid distributions in HUVECs using the SFN system at a 100-nm step size, enabling direct interrogation of pseudopodial lipid metabolism and elucidation of structure-biochemistry relationships.

Initial mapping at 2930 cm⁻¹ reconstructed cellular morphology. SFN enabled unambiguous delineation of the plasma membrane boundary, internal subcellular architecture, and fine extracellular edge features, demonstrating superior contrast and resolution (Fig. 5a). Specifically, high-resolution Raman images resolved four spatially distinct features (F1-F4) within the pseudopodium: a prominent, high-intensity focal spot (F1); a well-defined tubular structure extending longitudinally through the pseudopodium interior (F2); a granule-associated configuration at an intermediate position near the plasma membrane, demarcating the membrane boundary with multiple inward-facing protrusions (F3); and a distinct granular feature at the distal tip (F4). In stark contrast, conventional CRM acquired under identical conditions yielded only a diffuse, low-contrast signal at the F1 location and failed to resolve F2, F3, or F4, revealing no discernible boundaries or internal structures (Fig. 5b).

**Fig. 5.**
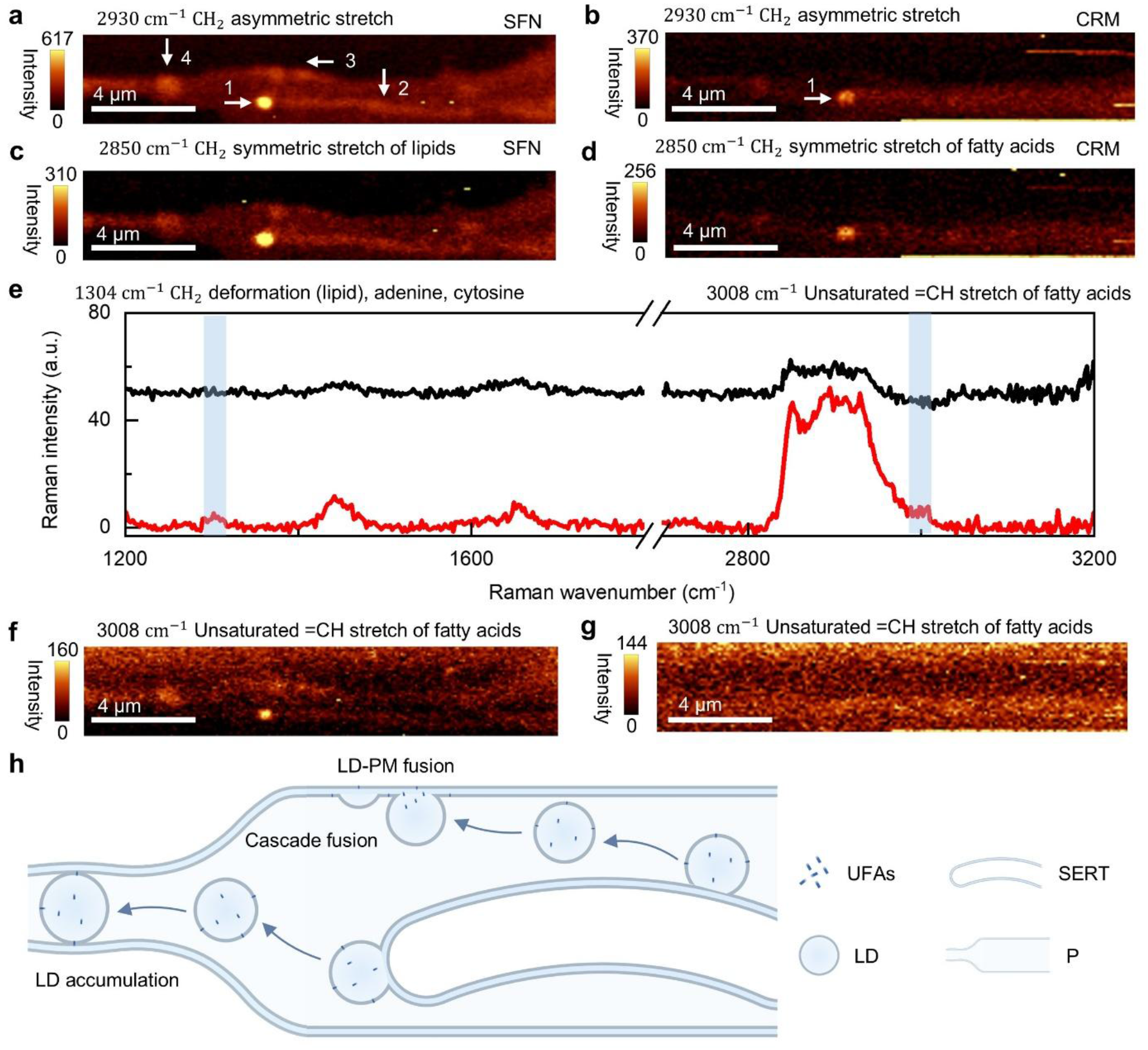
Nanoscale Raman mapping of lipid metabolism in HUVEC pseudopodia. **a**, SFN Raman image of a pseudopodium acquired at 2930 cm⁻¹, resolving four distinct features (F1–F4). **b**, Conventional CRM Raman image of the same pseudopodium acquired at 2930 cm⁻¹. **c**, SFN Raman image of a pseudopodium acquired at 2850 cm⁻¹, highlighting enhanced contrast and spatial definition of lipid-rich domains under SFN. **d**, Conventional CRM Raman image of the same pseudopodium acquired at 2850 cm⁻¹. **e**, Representative Raman spectra extracted from F1 in **a**, comparing spectral sensitivity between SFN and CRM. SFN resolves additional vibrational bands at 1304 cm⁻¹ and 3008 cm⁻¹, both absent in the CRM spectrum. **f**, SFN Raman image of the same pseudopodium acquired at 3008 cm⁻¹, revealing heterogeneous spatial enrichment of UFAs within distinct subcompartments. **g**, Conventional CRM Raman image of the same pseudopodium acquired at 3008 cm⁻¹. **h**, Schematic model of pseudopodial lipid trafficking pathways. Annotations: LD, lipid droplet; PM, plasma membrane; UFAs, unsaturated fatty acids; P, pseudopodium; SERT, smooth endoplasmic reticulum tubule. Acquisition parameters: 100 nm step size, 20 mW laser power, 1.0 s integration time.

Furthermore, we mapped the 2850 cm⁻¹ band to probe lipid distribution. As shown in Fig. 5c, SFN resolved three granular structures with distinct lipid signatures (F1, F3, and F4) and a continuous tubular element (F2). Morphologically, F2 displayed a membrane-like contour consistent with smooth endoplasmic reticulum tubule (SERT), and F1, physically contiguous with F2, corresponded to nascent LDs budding from these tubules. F3, positioned at the plasma membrane boundary with inward-facing protrusions, represented LDs engaged in membrane fusion, whereas F4, a discrete non-fused entity at the distal tip, corresponded to LDs accumulated for storage. In contrast, CRM again failed to resolve these features, yielding only a diffuse signal at the F1 location (Fig. 5d).

To rigorously evaluate spectral sensitivity, we performed detailed comparative analysis of Raman spectra acquired from region F1. Multiple additional vibrational modes were resolved at 1304 cm⁻¹ and 3008 cm⁻¹ (unsaturated =CH stretch of lipids)^25^, both of which remained undetectable under identical conditions using conventional CRM (Fig. 5e). The 3008 cm⁻¹ band, uniquely resolved by SFN, was spatially mapped to reveal its distribution across multiple subcellular compartments within the pseudopodium, including tubular structures and granular domains (Fig. 5f), providing direct spectroscopic evidence for heterogeneous enrichment of unsaturated fatty acids (UFAs) in functionally distinct pseudopodial microdomains. By contrast, conventional CRM yielded no discernible spectral or imaging information from the pseudopodium (Fig. 5g). This enhanced sensitivity and imaging capability is enabled by the synergistic action of PRE and 3D-SFE, consistent with the detection and resolution gains observed in the above experiments.

Notably, F3 exhibited features suggestive of cascading membrane fusion events. This phenomenon is likely attributable to three synergistic factors. First, fusion-promoting proteins are locally enriched at this site, providing the molecular machinery for LD-plasma membrane (LD-PM) fusion^29^. Second, the high local concentration of unsaturated fatty acids reduces membrane surface tension, thereby lowering the energy barrier for fusion^30,31^. Third, spatial confinement within the narrow pseudopodium increases the probability of contact between LDs and the plasma membrane^32,33^. Together, these factors may account for the high efficiency of LD-PM fusion at F3, ensuring a sufficient lipid supply for rapid pseudopodial extension.

Collectively, this spatial arrangement points to a model in which LDs synthesized at SERTs are delivered to two distinct endpoints: fusion with the plasma membrane at F3, or storage at F4. SFN- based mapping directly visualized all key stages of both pathways (Fig. 5h). This submicron-scale metabolic architecture establishes a direct link between nanoscale structural imaging and biochemically specific functional mapping, a level of insight previously beyond the reach of label- free Raman microscopy.

## Discussion

SFN achieves a lateral resolution of approximately 40 nm in label-free Raman imaging of both inorganic samples and biological specimens including silicon nanostructures, intact cells, and tissue sections through fundamental signal purification. Unlike conventional super-resolution approaches that rely primarily on signal amplification, SFN delivers effective super-resolution predominantly via background suppression: a robust, physically grounded signal-purification strategy that enables nanoscale, chemically specific imaging on standard confocal Raman microscope platforms without hardware modification or nonlinear excitation. Using SFN, we successfully resolved subcellular organelles and mapped pseudopodial lipid metabolism with molecular specificity. The achieved resolution not only substantially surpasses that of all existing label-free Raman super-resolution techniques but also approaches the performance levels attained by fluorescence-based super-resolution methods (Supplementary Table 1). Moreover, SFN offers practical advantages: straightforward implementation on commercial confocal microscopes, full compatibility with off-the-shelf optical components, and seamless integration with complementary modalities including fluorescence confocal microscopy, super-resolution fluorescence microscopy, and nonlinear Raman imaging.

SFN must be clearly distinguished from conventional microsphere-assisted microscopy (MAM). Although both techniques involve placing a microsphere on the specimen surface, their underlying physical principles are fundamentally different. MAM enhances resolution through photonic nanojets or evanescent-wave coupling and is compatible with both wide-field and confocal configurations. However, MAM microspheres, typically several to tens of micrometers in diameter, are subject to an intrinsic trade-off: resolution degrades with increasing sphere size. Additional limitations include an extremely restricted field of view, stringent requirements for microsphere quality and precise positioning, and spherical aberrations induced by the curved interface. In contrast, SFN repurposes the SMML as an integrated optical element that simultaneously compresses the excitation profile via photonic redistribution effect (PRE) and enables three-dimensional spatially filtering effect (3D-SFE) when combined with the confocal pinhole, bypassing reliance on nanojets or evanescent-wave coupling. As a result, SFN achieves high-resolution, high-contrast Raman imaging and highly sensitive Raman spectroscopy concurrently under standard excitation power and acquisition conditions without requiring elevated laser intensity. This operational advantage significantly reduces photothermal and photochemical damage to delicate biological specimens. Furthermore, SFN provides distinct practical benefits: a markedly enlarged field of view, compatibility with standard objective housings, and simplified handling, making it a robust, reproducible, and deployable platform.

Beyond its immediate technical impact, the geometric-optical framework underpinning SFN, specifically PRE and 3D-SFE, establishes a quantitative design foundation for SMML-assisted imaging and guides the rational engineering of next-generation SFN systems. More broadly, the SFN concept extends well beyond Raman microscopy. Because PRE and 3D-SFE are modality-agnostic physical effects acting independently on the excitation and collection pathways of any confocal system, the same hardware principle can be directly adapted to confocal fluorescence, photoluminescence, or stimulated Raman scattering (SRS) microscopy. In each case, the signal- purification paradigm offers a universal, hardware-native pathway to break the diffraction limit. Finally, we acknowledge current limitations of the SFN platform and outline concrete strategies for their mitigation: (1) Throughput and temporal resolution: A per-pixel dwell time of 0.5-1 s results in several hours to acquire a 30 μm × 30 μm super-resolution map. While the low phototoxicity of SFN permits prolonged imaging, this timescale remains incompatible with dynamic subcellular processes. Algorithmic acceleration including sparse sampling, compressed sensing, and deep learning-based reconstruction, is expected to reduce acquisition time by one to two orders of magnitude without compromising spatial fidelity. (2) Alignment and integration: Current SMML placement relies on manual micromanipulation, limiting reproducibility and scalability. Integrating the SMML directly into the objective housing via custom lens mounts or embedded microsphere elements, would eliminate manual alignment, enabling routine, operator- independent deployment.

## Methods

### SFN Instrumentation

The SFN system was implemented by integrating an SMML into a commercial confocal Raman microscope (alpha300R, WITec). All measurements were performed using a 63×/1.0 NA water-immersion objective (W Plan-Apochromat, Zeiss), and a 532 nm laser. K9 SMMLs (diameters: 300, 500, 800, and 1000 μm) and fused silica (FS) SMMLs were procured from Changzhou Runchang Optoelectronics (China). Polystyrene (PS) SMMLs were obtained from Yiyuan Bio (China). Unless otherwise specified, all data presented in the main figures were acquired under a fixed, optimized configuration using a 500 μm K9 SMML.

Raman spectra were dispersed by a 600 lines mm⁻¹ grating and detected with a thermoelectrically cooled CCD camera. For WFM and SIL imaging, the confocal pinhole and diffraction grating were removed, and direct optical imaging was performed using an integrated sCMOS camera.

### Sample Preparation

Four types of silicon substrates were employed: (i) A polished silicon wafer served as a standardized optical reference for Airy-disk characterization, quantitative Raman intensity calibration, and system alignment; (ii) A silicon nanostructure with regular square arrays was used to determine the effective magnification factor introduced by SMML coupling; (iii) Two silicon substrates featuring linear nanostructures, fabricated with nominal widths of 108 nm and 40 nm, were utilized to assess lateral resolution. Their precise dimensions were validated prior to Raman imaging via atomic force microscopy (AFM). All silicon substrates were rigorously cleaned by immersion in piranha solution (H₂SO₄:H₂O₂ = 7:3, v/v) for 30 min, followed by thorough rinsing with ultrapure water and drying under a stream of nitrogen.

Human umbilical vein endothelial cells (HUVECs) and human lung adenocarcinoma epithelial cells (A549) were cultured in DMEM supplemented with 10% fetal bovine serum (FBS) and 1% penicillin-streptomycin at 37 °C under 5% CO₂. For Raman imaging, cells were seeded on glass-bottom confocal dishes, washed three times with phosphate-buffered saline (PBS), fixed with 4% paraformaldehyde (PFA) for 15 min at room temperature, and rinsed again with PBS before measurement.

Mouse lung tissue sections (15 μm thickness) were prepared by fixing fresh tissue in 4% PFA at 4 °C for 24 h, cryoprotecting in optimal cutting temperature (OCT) compound, and sectioning using a cryostat. Sections were washed with PBS to remove residual OCT prior to Raman imaging.

### Magnification Calibration

The effective magnification imparted by SMML coupling was quantified using the silicon nanostructure with regular square arrays. First, a wide-field image of the nanostructure was acquired using the WFM mode. The physical pitch of the array was measured from this image using ImageJ and taken as the reference dimension. Second, an identical region was imaged under WFM mode with SMML coupling. The apparent pitch was measured analogously. The magnification factor was calculated as the ratio of the measured pitch with SMML to that without SMML.

To confirm consistency across modalities, the same nanostructure was imaged via Raman mapping at the Si-Si vibrational mode (520 cm⁻¹), using a step size of 300 nm, 0.5 s integration time, and excitation power of 20 mW. The magnification derived from Raman mapping was cross-validated against the WFM result.

### Airy Disk Characterization

Airy-disk measurements including signal intensity enhancement and FWHM were conducted on the polished Silicon wafer.

i. Signal enhancement: Wide-field images of the Airy disk were acquired at four incident laser powers in both WFM (0.1, 0.2, 0.3, and 0.4 mW) mode and SIL mode (0.1, 0.15, 0.2, and 0.3 mW). Integrated peak gray values within the first-order Airy bright ring were extracted using ImageJ. Signal enhancement was defined as the ratio of peak gray value with SMML to that without SMML.
ii. The diameter of the central Airy spot: Simultaneous illumination with 0.1 mW 532 nm laser and 0.1% white-light background was applied to the silicon wafer under both WFM and SIL modes. Airy-disk profiles were fitted with a Gaussian function.

### Raman Spectral Linearity and Background Assessment

To evaluate the linearity of SFN, Raman spectra of the polished silicon wafer were acquired using SFN configurations across a laser power series (0.25, 0.50, 1.0, 1.5, 3, 5, 10, 15, and 20 mW), with 0.5 s integration time per spectrum and 10 accumulations. After background subtraction, the peak intensity of the Si-Si stretching mode at 520 cm⁻¹ was plotted versus laser power to determine linearity (reported as R²). Background intensity was estimated as the mean signal in the Raman-silent region (1800-2700 cm⁻¹); noise was quantified as the standard deviation of that same region. **Raman Imaging Protocols**

- Silicon nanostructure imaging: Raman maps of the 108 nm square-array Si pattern were acquired using SFN and CRM. Acquisition parameters: 30 nm step size, 20 mW laser power, 1.0 s integration time. Bright-field reference images of the same region were acquired in WFM and SIL modes. Image contrast was quantified using the Michelson contrast formula: Cₘ = (Iₘₐₓ − Iₘᵢₙ)/(Iₘₐₓ + Iₘᵢₙ).
- Resolution validation on Silicon: Raman imaging of the 40 nm linear nanostructure Si pattern was performed under identical parameters to assess ultimate spatial resolution. Acquisition parameters: 30 nm step size, 20 mW laser power, 1.0 s integration time.
- Cellular imaging: Raman maps of A549 cells were acquired at step sizes of 200 nm (overview) and 30 nm (high-resolution), using 20 mW laser power and 1.0 s integration time. Vibrational bands mapped included 2930 cm⁻¹, 2850 cm⁻¹, 1445 cm⁻¹, 1339 cm⁻¹, 1304 cm⁻¹ and 1259 cm⁻¹. Parallel CRM imaging of identical regions served as controls.
- Tissue imaging: Mouse lung tissue sections were imaged at 200 nm and 30 nm step sizes (20 mW, 1.0 s/pixel), mapping 2930 cm⁻¹ and 1445 cm⁻¹. CRM control maps were acquired in parallel. Acquisition parameters: 20 mW laser power, 1.0 s integration time.
- Subcellular organelle imaging: High-resolution Raman maps of HUVECs were acquired using raster scanning with a 200 nm step size for whole-cell imaging and a 100 nm step size for subcellular structures, such as organelles and pseudopodia. Key biochemical bands included 2930 cm⁻¹, 2850 cm⁻¹, 3008 cm⁻¹, and 1670 cm⁻¹. Acquisition parameters: 20 mW laser power, 1.0 s integration time.

### Atomic Force Microscopy

All Silicon nanostructure topographies were characterized using a Bruker MultiMode 8-HR AFM operated in tapping mode under ambient conditions.

### Theoretical Modeling

Comprehensive derivations of the optical model, including PRE and 3D-SFE, are provided in Supplementary Notes 3-13.

## Data Analysis

All Raman images were processed using WITec Project 6.1 and ImageJ (NIH). Lateral resolution was determined by Gaussian fitting; the FWHM of the fitting was reported as the resolution metric.]

## Ethics declarations

Male BALB/c mice at 4-6 weeks old were purchased from Shanghai SLAC Laboratory Animal Co., Ltd. (License No. SCXK (Hu) 2019-0002). All animals were housed under specific pathogen-free (SPF) conditions and acclimatized for 5 days prior to experimentation. The study protocol was approved by the Institutional Animal Care and Use Committee (IACUC) of the School of Pharmaceutical Sciences, Fudan University (Approval No. 2020-09-SY-LJY-01). Human non-small cell lung carcinoma A549 cells and HUVECs were obtained from the Cell Bank of the Chinese Academy of Sciences. Cells were cultured according to the supplier’s recommendations and used within 10 passages.

## Data availability

The raw data that support the findings of this study are available from the corresponding author upon reasonable request.

## Acknowledgements

We acknowledge the Core Facilities of the School of Pharmaceutical Sciences at Fudan University for providing access to the Raman imaging platform. We thank the technical support team at Oxford Instruments (WITec) Shanghai for assistance with the confocal Raman microscope system.

## Author contributions

Hulie Zeng conceived and supervised the study and designed the experiments. Jiye Li developed the theoretical model, performed the theoretical calculations, conducted the Raman imaging experiments, and processed the data. Benma Lamu, Yuan Feng and Haiyan Zhu assisted with biological sample preparation. Shengwen Wang performed the atomic force microscopy (AFM) characterization. Jiye Li, Hulie Zeng and Zhiru Xu wrote the manuscript. All authors discussed the results, interpreted the data, and approved the final manuscript.

---

**Extended Data Fig. 1.**
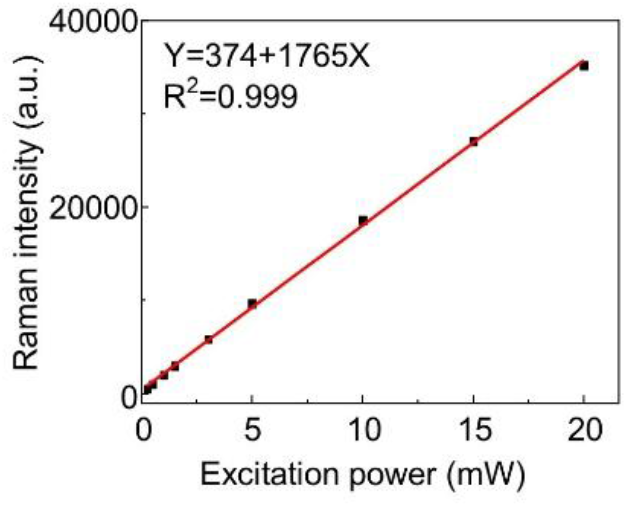
Linear response of the SFN system. Raman signal intensity of the silicon wafer at the 520 cm⁻¹ Si-Si mode as a function of excitation laser power. The data exhibit excellent linearity (R² > 0.999), confirming that the SFN system operates within the linear response regime without detector saturation or nonlinear effects. Acquisition parameters: 0.5 s integration time, 10 accumulations. Data are presented as mean ± s.d (N = 5).

**Extended Data Fig. 2.**
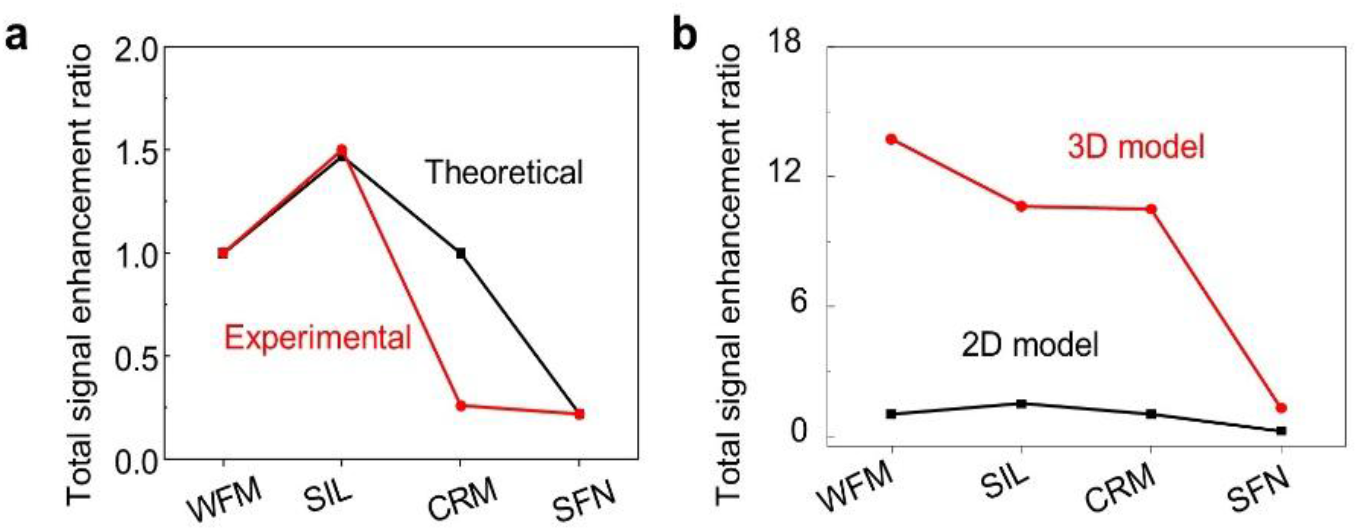
Total signal enhancement ratios across configurations. a, Theoretical versus experimental ratios for WFM, SIL, CRM, and SFN, normalized to 2D WFM. **b**, Theoretical total signal enhancement ratio for 2D and 3D models across WFM, CRM, SIL and SFN configurations, normalized to 2D WFM.

**Extended Data Fig. 3.**
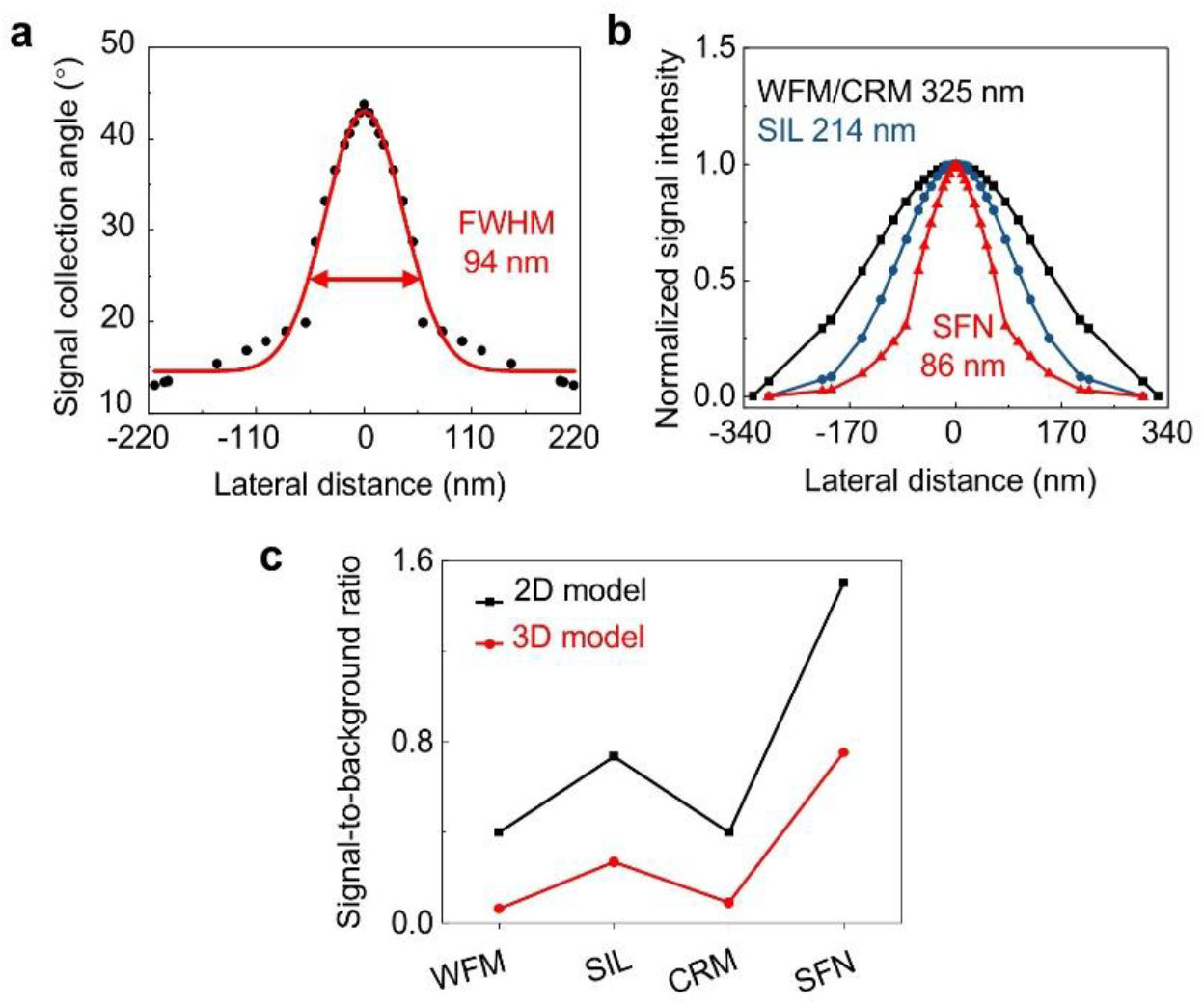
Theoretical analysis of resolution and signal-to-background ratio of SFN. **a**, Theoretical lateral resolution limited by signal collection alone. **b**, Theoretical lateral resolution incorporating both Raman signal collection and laser excitation. **c**, Theoretical signal-to-background ratio for 2D and 3D models across WFM, SIL, CRM and SFN configurations.

**Extended Data Fig. 4.**
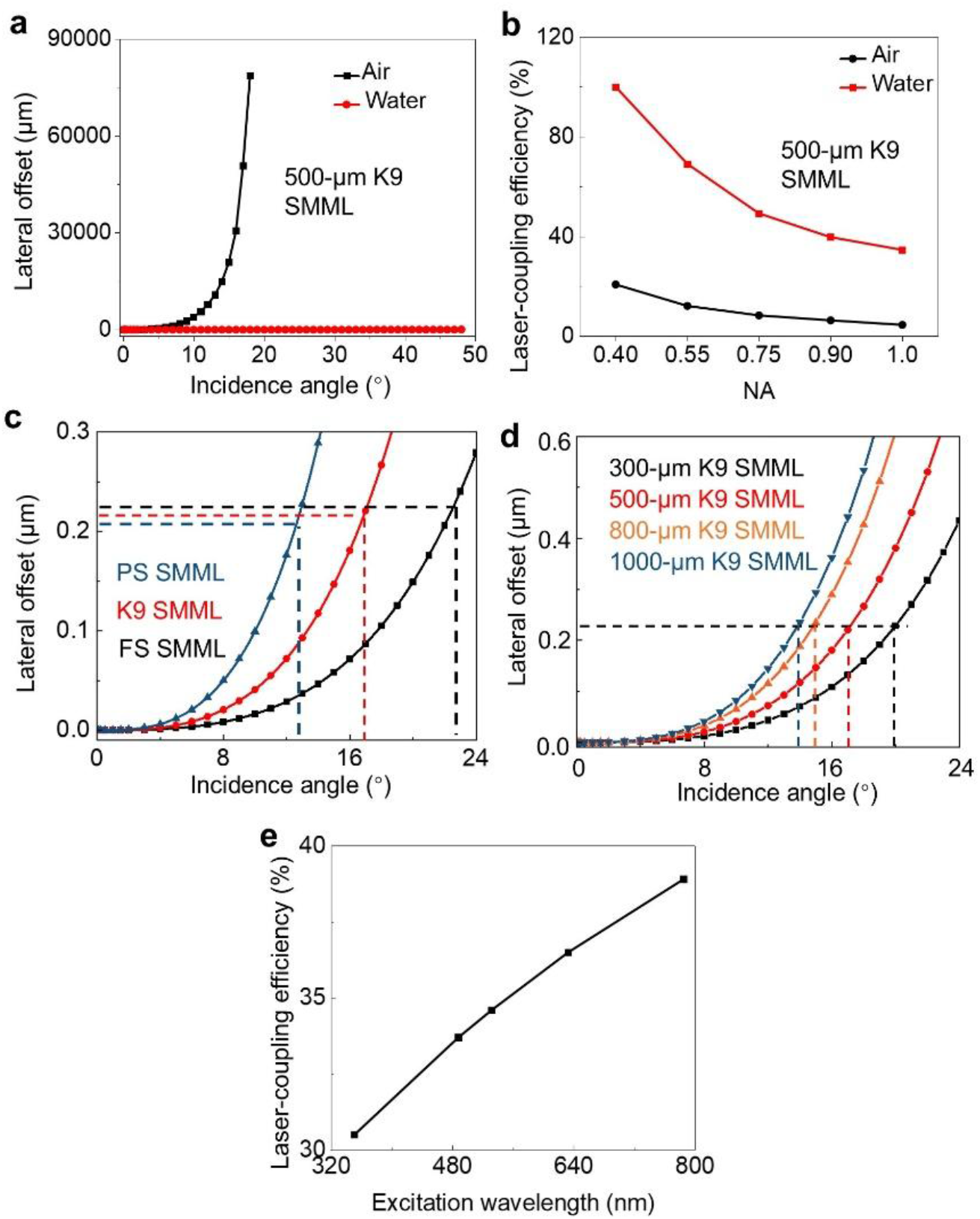
Laser excitation-coupling efficiency under different optical conditions. **a**, Theoretical laser excitation-coupling efficiency as a function of incidence angle adopting K9 SMMLs in air or water. **b**, Dependence of the laser excitation-coupling efficiency on objective NA adopting 500-μm K9 SMML. **c**, Calculated lateral offset as a function of incidence angle for different refractive indices of SMMLs. **d**, Theoretical laser excitation-coupling efficiency as a function of incidence angle for different diameters of K9 SMMLs. **e**, Dependence of laser excitation-coupling efficiency on excitation wavelength utilizing 500- μm K9 SMML.

**Extended Data Fig. 5.**
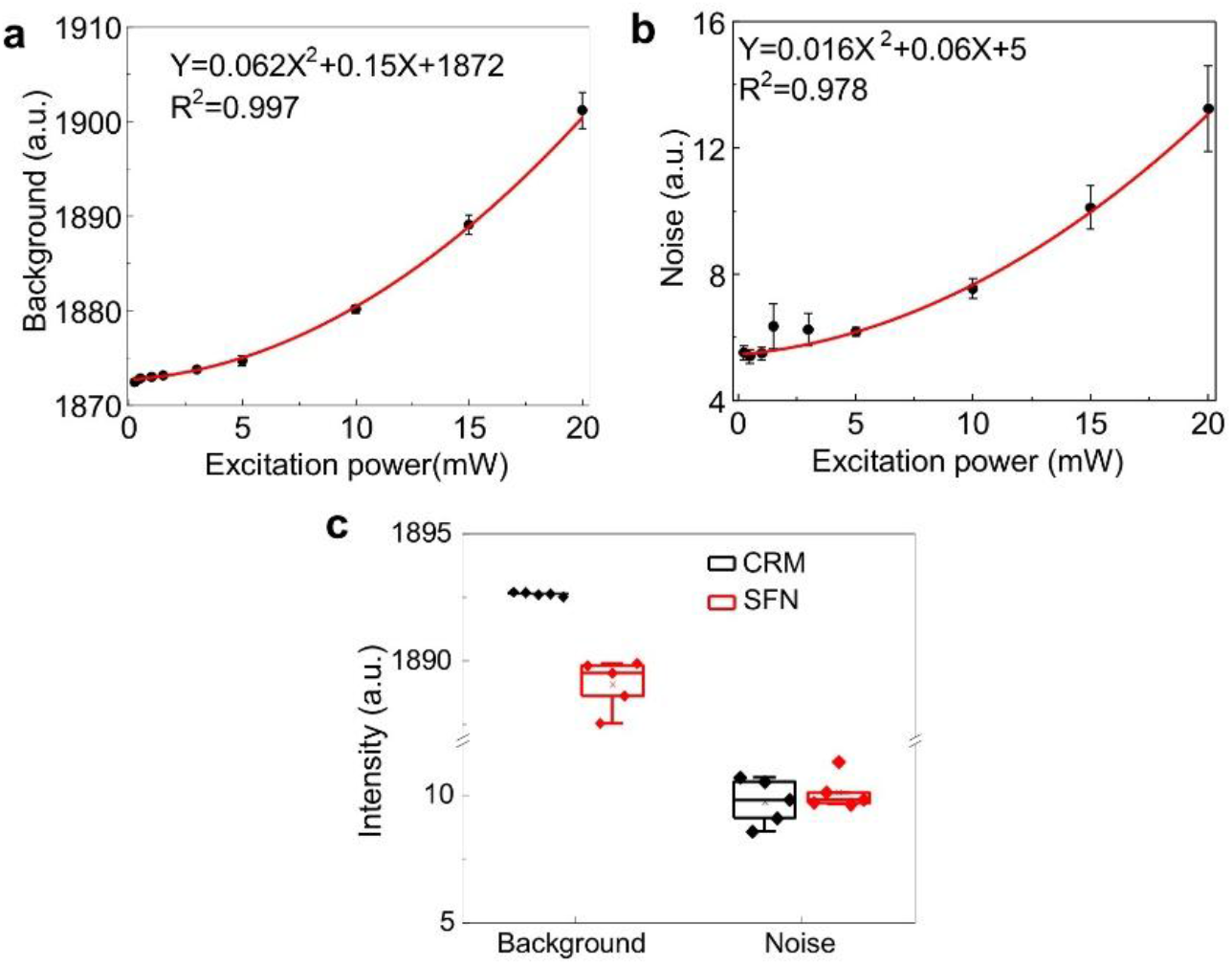
Spectral characterization of SFN system. **a**, Background intensity as a function of laser power, showing quadratic scaling. **b**, Noise level as a function of laser power, also showing quadratic scaling. **c**, Comparison of Raman spectra acquired with conventional CRM (10 mW) and SFN (15 mW). Data are presented as mean ± s.d (N = 5).

**Extended Data Fig. 6.**
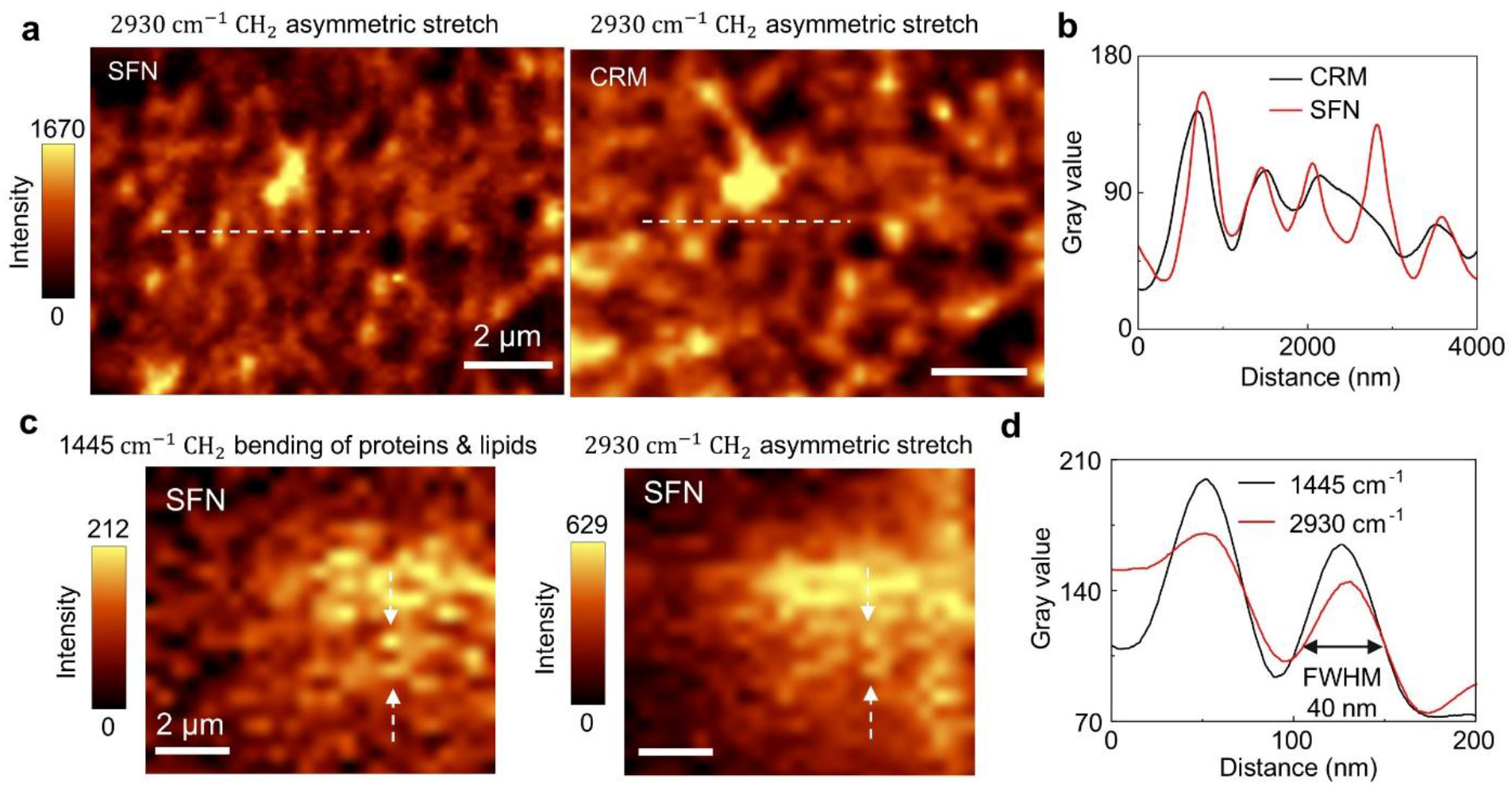
Super-resolution imaging of mouse lung tissue acquired by SFN. **a**, Raman images of lung tissue at 2930 cm⁻¹ acquired by SFN (left) and conventional CRM (right), with corresponding lateral intensity profiles along the dashed lines. Acquisition parameters: 200 nm step size, 20 mW laser power, 1.0 s integration time per pixel. **b**, Intensity profiles along white dashed line in **a**. **c**, Multispectral SFN Raman images of lung tissue mapped at 1445 and 2930 cm⁻¹. Acquisition parameters: 30 nm step size, 20 mW laser power, 1.0 s integration time per pixel. **d**, Intensity profiles between two white dashed arrows in **c**.

**Extended Data Fig. 7.**
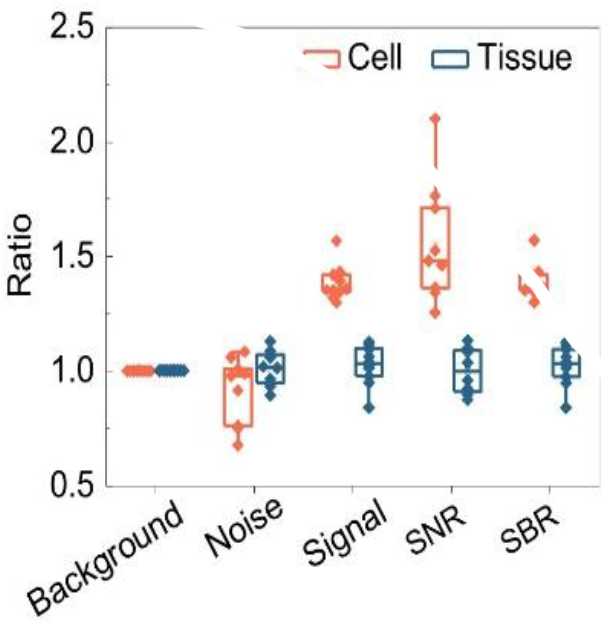
Quantitative comparison of Raman signal performance in cells and tissues utilizing SFN system. Comparing normalized signal, background, noise, SNR and SBR in cells (red) and tissues (blue). All values are normalized according to the signals of CRM system. Data are presented as mean ± s.d (N = 10).

