## Supplementary Information for "Spatial-filtering nanoscopy for 40-nm label-free Raman imaging"

Jiye Li (李吉业)[1,2], Benma Lamu (本玛拉姆)[1], Yuan Feng (冯瑗)[4], Shengwen Wang (王圣文)[5], Haiyan Zhu (朱海燕)[4], Zhiru Xu (徐智儒)[2,3], Hulie Zeng (曾湖烈)[1]

[1]Department of Pharmaceutical Analysis, School of Pharmaceutical Sciences, Key Laboratory of Smart Drug Delivery, Ministry of Education, Fudan University, Shanghai 201203, China;

[2]China State Institute of Pharmaceutical Industry, Shanghai 201203, China;

[3]National Key Laboratory of Lead Druggability, Shanghai Institute of Pharmaceutical Industry, Shanghai Professional and Technical Service Center for Biological Material Druggability Evaluation, Shanghai 200437, China;

[4]Department of Biological Medicines, School of Pharmaceutical Sciences, Fudan University, Shanghai 201203, China;

[5]Institute of Molecular Medicine and Shanghai Key Laboratory for Nucleic Acid Chemistry and Nanomedicine, Renji Hospital, School of Medicine, Shanghai Jiao Tong University, Shanghai 200127, China;

### Supplementary Notes

#### 1. Geometrical-optics derivation of the magnification of the SMML

Because the dimensions of SMML significantly exceed the optical wavelength, diffraction and interference effects are negligible; thus, its imaging behaviour can be rigorously modelled within the framework of geometrical optics. Accordingly, the SMML may be treated as a thick lens, whose focal length $f$ satisfies the standard thick-lens formula:

$$\frac{1}{f}=\left(\frac{n-1}{R_1}\right)+\left(\frac{n-1}{R_2}\right)-\left(\frac{(n-1)^2 d}{nR_1R_2}\right) \qquad (S1-1)$$

where $R_1$ and $R_2$ denote the radius of the first and second surfaces, $d$ is the lens thickness (equal to the SMML diameter), and $n$ is the relative refractive index defined as:

$$n=\frac{n_{\mathrm{SMML}}}{n_{\mathrm{A}}} \qquad (S1-2)$$

where $n_{\mathrm{SMML}}$ represents the absolute refractive index of the SMML material and $n_{\mathrm{A}}$ is that of the ambient medium. For a symmetric SMML, $R_1 = R_2 = R = d/2$, reducing Eq. (S1-1) to:

$$\frac{1}{f}=\frac{2(n-1)}{nR} \qquad (S1-3)$$

This focal length also obeys the equation:

$$\frac{1}{f}=\frac{1}{u}+\frac{1}{v} \qquad (S1-4)$$

where $u$ and $v$ are the object and image distances, respectively. Given that the SMML is placed in direct contact with the sample surface, $u = R$. Substituting into Eq. (S1-3) and (S1-4) yields:

$$v=\frac{nR}{n-2} \qquad (S1-5)$$

When $n < 2$, $v$ is negative, indicating formation of a virtual image. The corresponding lateral magnification $M$ is given by:

$$M = -\frac{v}{u} \qquad (S1-6)$$

Substituting Eq. (S1-5) and (S1-6) yields

$$M = \frac{n}{2-n} \qquad (S1-7)$$

Eq. (S1-7) aligns quantitatively with prior reports[1].

To experimentally validate Eq. (S1-7), we first evaluated K9 SMMLs of varying diameters (300, 500, 800, and 1000 μm) under both SMML-coupled wide-field microscopy (SIL) and SFN. All lenses resolved the silicon nanostructure with high fidelity (Supplementary Fig. S1a,b). Measured magnifications agreed closely with theoretical predictions across all diameters (Supplementary Fig. S1c), confirming that magnification is effectively independent of SMML diameter.

Next, we systematically varied the relative refractive index $n$ using 500-μm-diameter SMML fabricated from fused silica (FS), K9 glass, and polystyrene (PS). In both SIL and SFN configurations, all three materials successfully resolved the nanostructure (Supplementary Fig. S2a,b). Experimentally determined magnifications matched theoretical values well and increased monotonically with $n$ (Supplementary Fig. S2c), corroborating that magnification is predominantly governed by the $n$ of SMMLs.

Although all SMMLs achieved nanostructure imaging, image quality was slightly degraded for the 300-μm K9, 500-μm FS, and 500-μm PS SMMLs, evidenced by reduced edge sharpness. This degradation is likely attributable to fabrication imperfections, with local surface roughness further scattering incident light and thereby reducing the effective numerical aperture. Furthermore, the PS SMML exhibited stronger chromatic dispersion in SIL imaging, a consequence of its higher material dispersion (lower Abbe number) relative to the FS and K9 SMMLs.

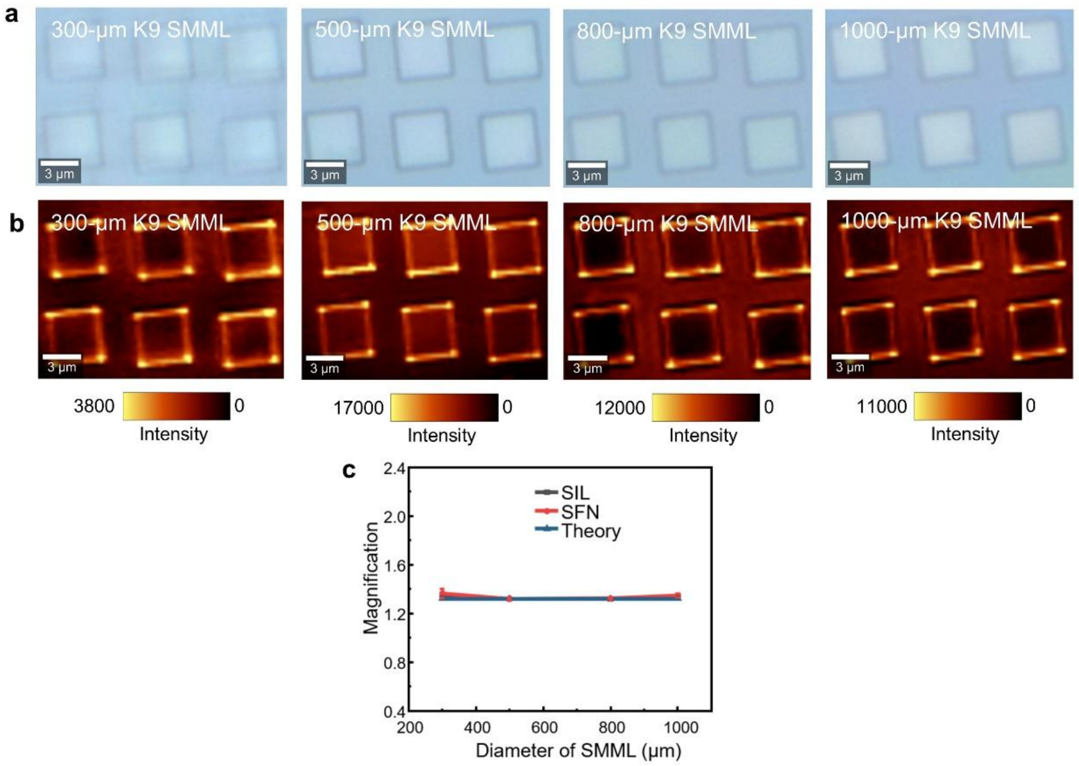

**Supplementary Fig. S1 | Influence of SMMLs diameter on magnification. a**, Bright-field images of the silicon nanostructure acquired using K9 SMMLs of different diameters under SIL mode. **b**, Raman images of the same silicon nanostructure acquired using K9 SMMLs of different diameters under SFN mode. **c**, Comparison of theoretical and experimental magnifications as a function of the K9 SMML diameter. Acquisition parameters of all Raman imaging: 300 nm step size, 520 cm$^{-1}$ Si-Si mode, 0.5 s integration time and 20 mW laser power. Data are presented as mean values ± s.d. over the field of view (FOV).

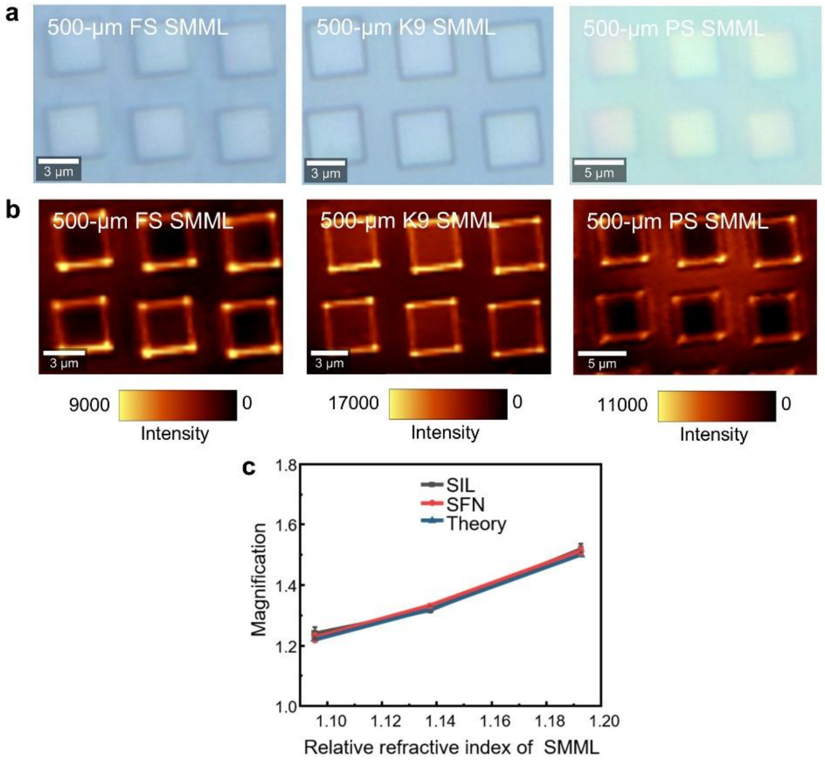

**Supplementary Fig. S2 | Influence of refractive index of SMMLs on magnification. a**, Bright-field images of the silicon nanostructure acquired using SMMLs with different refractive indices under SIL mode. **b**, Raman images of the same silicon nanostructure acquired using SMMLs with different refractive indices under SFN mode. **c**, Comparison of theoretical and experimental magnifications as a function of the relative refractive index of the SMMLs. Acquisition parameters of all Raman imaging: 300 nm step size, 520 $cm^{-1}$ Si-Si mode, 0.5 s integration time and 20 mW laser power. Data are presented as mean values ± s.d. over the field of view (FOV).

#### 2. Linear-response verification of WFM and SIL by Airy disk

For a linear optical system, the measured signal intensity is expected to scale linearly with incident excitation power, provided that detector saturation, nonlinear absorption, thermal lensing, or other nonlinear photophysical effects are absent. The Airy disk represents the spatial distribution of signal intensity and thus serves as a natural probe for verifying this linearity. A polished silicon wafer was selected as the calibration substrate. Its atomically flat surface minimizes axial background contributions and provides a well-defined, specular planar interface ideal for quantitative intensity calibration. Using a 532 nm laser, we generated Airy disks on the wafer surface via both WFM and SIL configurations (Supplementary Fig. S3). However, the central Airy spot consistently suffered from detector saturation even at the lowest excitation power (0.10 mW), rendering it unsuitable for quantitative analysis (Fig. 2b). In contrast, the first-order Airy bright ring (termed Airy ring) remained unsaturated and exhibited stable, reproducible contrast. For quantitative analysis, we extracted the peak gray value at the brightest position along the Airy ring. We first validated linearity in the WFM configuration. Airy disks were acquired at incident powers of 0.10, 0.20, 0.30, and 0.40 mW (Supplementary Fig. S4). The peak gray value of the Airy ring was plotted against incident power, yielding an excellent linear fit ($R^2 = 0.996$), confirming that WFM operates linearly over this power range. Having established linearity in WFM, we extended the same measurement to the SIL configuration. Identical measurements were performed under the SIL configuration at powers of 0.10, 0.15, 0.20, and 0.30 mW (Supplementary Fig. S5). Again, the peak gray value of the Airy ring varied linearly with excitation power, with $R^2 = 0.983$, demonstrating that SIL maintains linear response under our experimental conditions.

Notably, both linear fits exhibited nonzero y-intercepts. This offset arises from additive background contributions, primarily dark current of the camera sensor and stray light within the optical path, both independent of laser power. Such constant offsets do not invalidate the linear relationship; rather, they represent a well-characterized baseline that can be subtracted during quantitative analysis. Their presence is expected and fully consistent with standard photometric calibration practice.

Although the power range used for the above verification is relatively narrow (0.10–0.30 mW), it is sufficient to establish the linearity of the optical configuration. Within a linear system, the transfer function is an intrinsic property of the optical path, independent of excitation power, provided that no saturation or nonlinear effects are triggered. The absence of such effects within the tested range therefore implies that linearity holds across the broader power regime. To further corroborate this inference, we independently confirmed the linear response of the SFN configuration over a substantially wider power range (0.25–20 mW) by measuring the Raman signal intensity of a silicon wafer as a function of laser power (Extended Data Fig. 1, $R^2 > 0.999$). Since SFN shares the identical excitation and collection optics with SIL (with the additional confocal pinhole), this wide-range linearity serves as strong evidence that the SIL

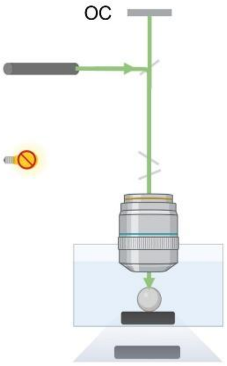

**Supplementary Fig. S3 | Schematic of the Laser illumination modes in SIL.**

configuration likewise remains linear under higher excitation powers.

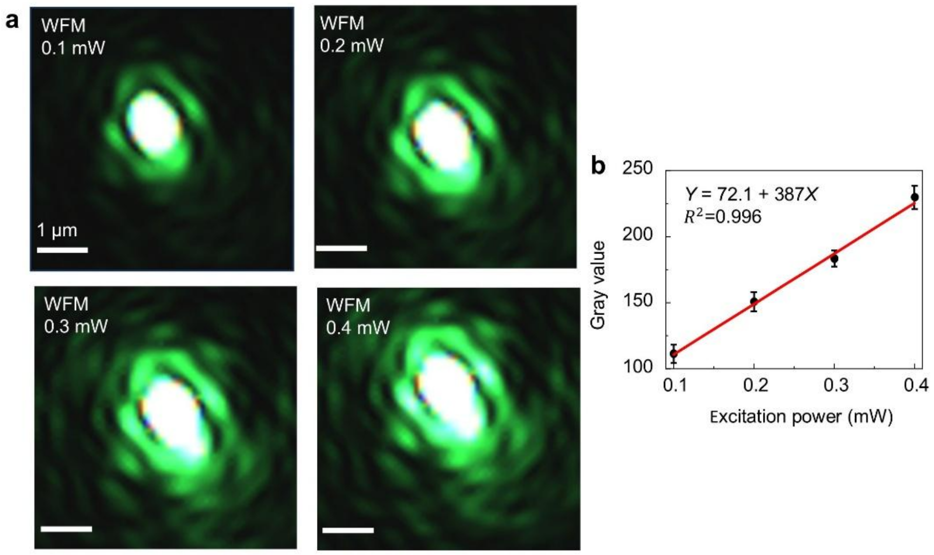

**Supplementary Fig. S4 | Dependence of intensity of Airy-ring on excitation power in WFM mode. a**, Airy disk images acquired under WFM mode at excitation powers of 0.10, 0.20, 0.30 and 0.40 mW. **b**, Linear dependence of the gray value of Airy-ring on excitation power in WFM. Data are presented as mean ± s.d. (N = 5).

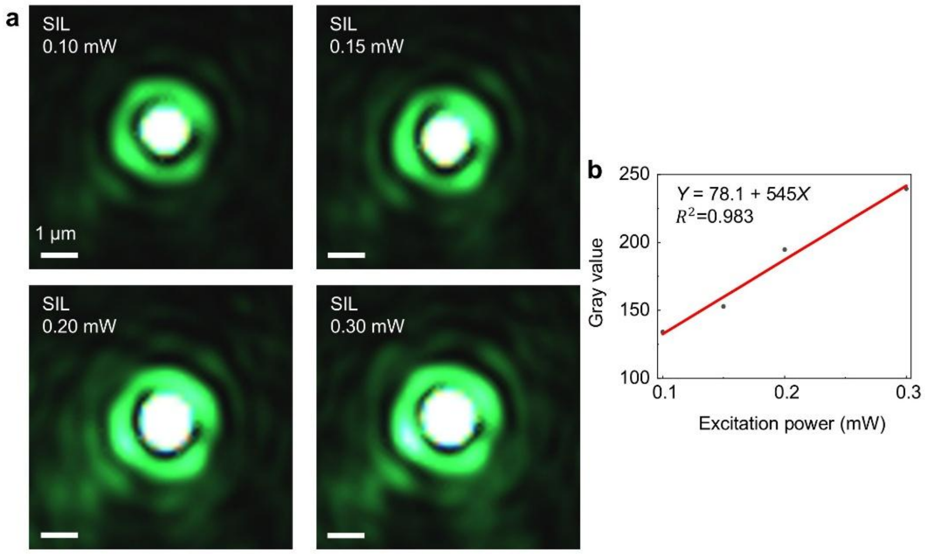

**Supplementary Fig. S5 | Dependence of intensity of Airy-ring on excitation power in SIL mode. a**, Images of Airy disk using SIL at excitation powers of 0.10, 0.15, 0.20 and 0.30 mW. The diameter of K9 SMML was 500 μm. **b**, Linear dependence of the Airy-ring gray value on excitation power in SIL. Scale bars were calibrated according to the measured magnification.

#### 3. Theoretical model of SMML-induced signal enhancement

We developed a quantitative optical model under the experimentally validated linear-response regime to describe the SMML-induced signal enhancement. This model provides a general theoretical framework for evaluating the signal enhancement capability of SMMLs across different imaging configurations. The total signal-enhancement ratio ($E_{\text{total}}$) can be factorized into excitation and collection terms:

$$E_{\text{total}} = E_{\text{exc}} \times E_{\text{col}} \qquad (S3-1)$$

where $E_{\text{exc}}$ quantifies the SMML-induced enhancement in effective excitation intensity, and $E_{\text{col}}$ represents the corresponding improvement in Raman signal collection efficiency. The excitation enhancement further decomposes into two physically distinct contributions: laser coupling efficiency $\eta_{\text{c}}$ and focusing gain $G_{\text{f}}$.

$$E_{\text{exc}} = \eta_{\text{c}} \times G_{\text{f}} \qquad (S3-2)$$

Where, $\eta_{\text{c}}$ denotes the fraction of incident laser power that remains collimated and effectively focusable after transmission through the SMML, while $G_{\text{f}}$ captures the intensity gain arising from the increased effective numerical aperture conferred by the SMML. We now derive each of these two components separately.

$\eta_{\text{c}}$ reflects the finite angular acceptance of the SMML for the objective-delivered excitation beam. Incident rays exceeding the critical acceptance angle of SMML undergo significant refraction and lateral displacement, reducing their contribution to the focused spot. Under a first-order geometric-optics approximation, the coupling efficiency is approximated as:

$$\eta_{\mathrm{c}} = \frac{\theta_{\mathrm{eff}}}{\theta} \qquad (S3-3)$$

where $\theta$ is the incidence angle of the excitation cone delivered by the objective in the conventional microscope, and $\theta_{\mathrm{eff}}$ is the effective incidence angle subtended by the SMML.

$G_{\mathrm{f}}$ originates from the SMML-induced increase in effective numerical aperture. Upon coupling into the higher-refractive-index SMML medium, the excitation beam converges more tightly, compressing the focal volume and thereby increasing local excitation intensity. Under the experimentally confirmed linear-response condition, this intensity enhancement directly scales the generated Raman signal. For an ideal optical system, $G_{\mathrm{f}}$ is given by the square of the NA ratio:

$$G_{\mathrm{f}} = \left(\frac{\mathrm{NA}_{\mathrm{eff}}}{\mathrm{NA}}\right)^2 \qquad (S3-4)$$

where NA is the numerical aperture of the original objective lens, and $\mathrm{NA}_{\mathrm{eff}}$ is the effective numerical aperture of the SMML-integrated objective system.

$E_{\mathrm{col}}$ quantifies the improved fraction of emitted photons captured by the detection objective. This arises from the ability of the SMML to broaden the signal collection angle, effectively increasing the maximum collection half-angle from $\alpha$ (objective) to $\alpha_{\mathrm{eff}}$ (in SMML-integrated the microscope system). $E_{\mathrm{col}}$ is expressed as:

$$E_{\mathrm{col}} = \frac{\alpha_{\mathrm{eff}}}{\alpha} \qquad (S3-5)$$

where $\alpha$ and $\alpha_{\mathrm{eff}}$ denote the signal collection half-angles before and after SMML integration, respectively.

#### 4. Determination of focusing gain

To characterize the saturated central Airy spot with quantitative fidelity, we implemented a dual-illumination strategy: simultaneous 0.1% white-light illumination (to elevate the background intensity level) and 0.1 mW 532 nm laser excitation (Supplementary Fig. S6). The white-light component raises the baseline signal uniformly across the field of view, thereby reducing the peak-to-background contrast of the central Airy spot. This avoids detector saturation at the focal centre while preserving the full spatial profile including the peak intensity within the linear dynamic range of the camera, enabling accurate, quantitative characterization of the central Airy spot.

To compute $G_f$ via Eq. (S3-4), the $NA_{eff}$ of the SIL system must first be determined experimentally. We extracted $NA_{eff}$ directly from the measured FWHM of the intensity distribution of central Airy spot, specifically its major-axis diameter $d_{Airy}$, using the diffraction-limited relation:

$$d_{\mathrm{Airy}} = \frac{1.22\lambda}{NA} \qquad (S4-1)$$

We validated this calibration approach using the WFM configuration as a reference. Under identical imaging conditions, the major-axis diameter of the central Airy spot was measured as 644.0 ± 35.3 nm, yielding an experimental $NA_{eff}$ of 1.01 ± 0.05. This value agrees with the manufacturer-specified NA of 1.0, confirming the reliability and accuracy of the NA determination method.

Applying the same procedure to the SIL configuration, the major-axis diameter of the central Airy spot was determined as 429.6 ± 9.2 nm, corresponding to $NA_{eff}$ of 1.51 ± 0.03. Substituting NA = 1.01 (WFM reference) and $NA_{eff}$ =1.51 into Eq. (S3-4) yields $G_f = (NA_{eff}/NA)^2 \approx 2.24$.

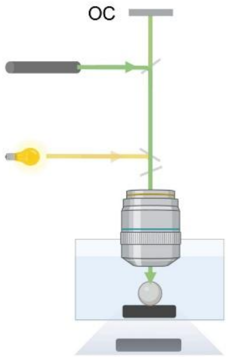

**Supplementary Fig. S6 | Schematic of the dual-source illumination modes for SIL.**

#### 5. Determination of laser-coupling efficiency

To quantify the fraction of incident laser power that remains collimated and effectively focusable after transmission through SMML, we modelled $\eta_c$ using geometric optics. The excitation objective employed is a 63×, NA = 1.0 water-immersion lens; under these conditions, the maximum incidence angle of the objective-delivered beam $\theta$ is 48.6°.

The effective angular acceptance of the SMML is governed by two sequential refractions: first, at the water-SMML interface (bottom spherical surface); second, at the SMML-water interface (top spherical surface) (Supplementary Fig. S7a,b). Both interfaces obey Snell's law. As the incident angle increases beyond a critical value, refracted rays undergo excessive lateral displacement and fail to converge within the diffraction-limited focal volume, and thus cannot be effectively focused. The limiting case corresponds to rays whose refracted trajectories just encompass the experimentally measured radius of the central Airy spot ($r_{\mathrm{Airy}}$). From the SIL-measured major-axis diameter of 429.6 ± 9.2 nm, $r_{\mathrm{Airy}}$ is approximately 215 nm. Using our self-consistent geometric-optics model, which incorporates SMML diameter, material refractive index, and immersion medium, we calculated the maximum input angle yielding a focused spot no larger than $r_{\mathrm{Airy}}$. This yields a $\theta_{\mathrm{eff}}$ of 16.8°.

Substituting $\theta = 48.6°$ and $\theta_{\mathrm{eff}} = 16.8°$ into Eq. (S3-3) gives: $\eta_c$ is approximately 0.35. Thus, approximately 35% of the incident laser power is coupled into focus-supporting modes within SMML; the remainder is lost primarily through angular rejection (i.e., refraction-induced beam deviation beyond the system's focusing tolerance) at SMML interfaces.

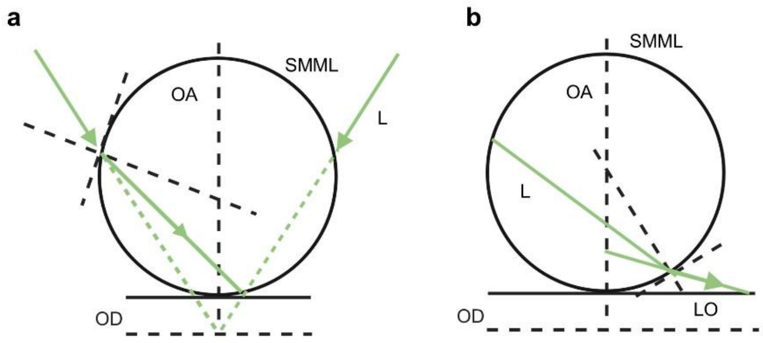

**Supplementary Fig. S7 | Geometrical optics model of laser coupling in SIL. a**, Geometrical-optics model of laser refraction entering the SMML. **b,** Geometrical-optics model of laser refraction exiting the SMML. L, laser ray; LO, lateral offset; OD, objective distance; OA, optical axis. Calculations were performed under experimental conditions with a 500-μm K9 SMML coupled to a 63×/1.0 NA water-immersion objective and 532 nm excitation.

**6. Determination of the signal-collection efficiency of SIL**

To quantify the fraction of signal captured by the detection objective after SMML integration, we determined the signal-collection efficiency $E_{col}$ using a ray-tracing geometric-optics model. For the bare 63×, NA = 1.0 water-immersion objective, the maximum collection half-angle $\alpha$ is approximately 48.6°. The objective width was calculated from its specified working distance (WD = 2.1 mm) and $\alpha$, yielding 4.77 mm (Supplementary Fig. S8a), consistent with manufacturer specifications.

To determine the $\alpha_{eff}$ in the SIL configuration, we constructed a self-consistent geometric-optics model (Supplementary Fig. S8b). In this model, the signal generated at the focal plane propagates through the SMML and undergoes two refractions at the SMML interfaces before being collected by the objective. Assuming an aperture half-angle of 90° (i.e., full aperture angle of 180°), we calculated the effective objective width required to collect the full angular range of the emitted signal. The calculation yielded an effective objective width of 2.90 mm, which is smaller than the physical objective width of 4.77 mm, confirming that the effective aperture half-angle reaches 90°. Substituting $\alpha$ = 48.6° and $\alpha_{eff}$ = 90° into Eq. (S3-5) gives: $E_{col} = \alpha_{eff} / \alpha \approx 1.85$. Thus, the SMML enhances the collected signal flux by a factor of approximately 1.85 relative to the bare objective.

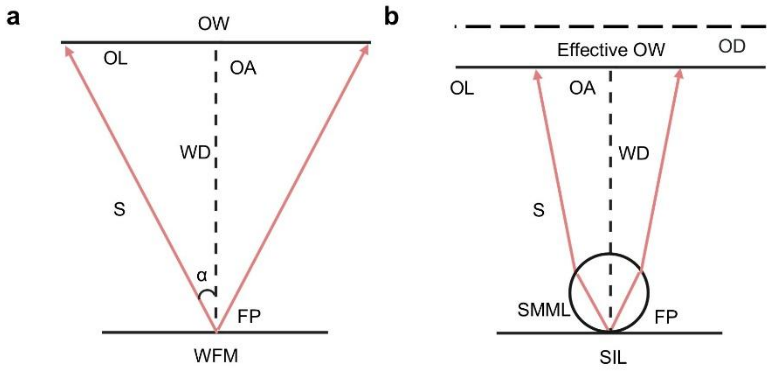

**Supplementary Fig. S8 | Geometrical optics model of Raman signal collection. a**, Geometrical optics model of the signal collection in WFM. **b**, Geometrical optics model of the signal collection in SIL. OW, objective width; OL, objective lens; OA, optical axis; WD, working distance; FP, focal point; α, aperture half-angle; OD, objective distance. Calculations were performed under experimental conditions with a 500-μm K9 SMML coupled to a 63×/1.0 NA objective.

#### 7. Qualitative geometric optics model for 3D-SFE by mapping SFN to CRM

To quantitatively assess the spatial filtering effect of the SFN system, we developed a geometric-optics model grounded in an equivalence principle. The core concept is to map the SFN configuration onto a well-characterized reference system, conventional CRM, whose axial PSF and depth-discrimination properties are rigorously established. The equivalence is established through the following procedure. In the SFN system, a signal ray emitted from the sample undergoes two refractions at the SMML interfaces and emerges from the SMML. The emerging ray is then extended backwards to intersect the optical axis. From this intersection point, we extract two key parameters: (i) the axial position $z$ along the optical axis, and (ii) the effective emission half-angle $\theta$ subtended at that point. These define an equivalent "virtual source" location and angular distribution, enabling direct comparison with known axial filtering response of CRM.

This equivalence framework yields three distinct analytical configurations:

(i) The focal-point model, applicable when the emission originates precisely at the focal point. Due to rotational symmetry about the optical axis, ray tracing need only be performed for a single meridional plane.

(ii) The lateral-offset model, used when the emitter is displaced laterally from the optical axis but remains at the focal plane. This breaks axial symmetry, necessitating full ray tracing.

(iii) The axial-offset model is adopted when the emission originates from a point that is displaced along the axial direction while remaining on the optical axis (i.e., zero lateral offset). The system retains mirror symmetry about the optical axis; thus, calculations remain confined to one meridional plane.

**8. Quantitative theoretical model for 3D-SFE**

Building upon the geometric equivalence framework established in Supplementary Note 7, we developed a quantitative theoretical model to compute the axial and lateral spatial filtering efficiency of the SFN system. To determine whether signal rays transformed by the SMML satisfy the confocal detection criterion, that is, whether they pass through the physical pinhole, we further constructed a ray-based geometric-optics model of conventional CRM, explicitly mapping axial source position and emission angle to detector transmission probability.

The PSF describes the intensity distribution of an ideal point source after imaging through a real optical system.

$$\mathrm{PSF_{total}} = \mathrm{PSF_{exc}} \times \mathrm{PSF_{col}} \qquad (S8-1)$$

where $\mathrm{PSF_{exc}}$ and $\mathrm{PSF_{col}}$ represent the contributions from excitation and collection, respectively. CRM employs a conjugate pinhole to reject out-of-focus light, rendering $\mathrm{PSF_{col}}$ the key factor governing axial signal transmission.

Empirical measurements show that confocal detection improves axial resolution by approximately 1.4×. Under the assumption of matched illuminatio and collection optics, the excitation and collection PSFs become comparable in axial extent, yielding the widely adopted approximation:

$$\mathrm{PSF_{exc}} \approx \mathrm{PSF_{col}} \qquad (S8-2)$$

Both PSFs are well approximated by Gaussian functions:

$$\mathrm{PSF} = e^{-\left(\frac{\rho^2}{2\delta^2}\right)} \qquad (S8-3)$$

where $\rho$ denotes either lateral *x-y* or axial *z* displacement from the focal point, and $\delta$ is the Gaussian standard deviation, related to the FWHM resolution $d_\mathrm{\rho}$ via:

$$\delta \approx \frac{d_\mathrm{\rho}}{2.355} \qquad (S8-4)$$

The FWHM resolution in WFM is given by classical diffraction theory:

$$d_\mathrm{lateral} = \frac{0.61 \times \lambda}{\mathrm{NA}} \qquad (S8-5)$$

$$d_\mathrm{axial} = \frac{2 \times n_\mathrm{A} \times \lambda}{\mathrm{NA}^2} \qquad (S8-6)$$

where $n_\mathrm{A}$ is the refractive index of the ambient medium (water, $n_\mathrm{A}$= 1.33). Within the CRM framework, the axial angular distribution of signal collection can be expressed as:

$$\Omega_{\mathrm{col}} \approx \alpha \times \mathrm{PSF}_{\mathrm{col}} \approx \alpha \times e^{-\left(\frac{z^2}{2\delta^2}\right)} \qquad (S8-7)$$

where $\alpha$ is the aperture half-angle of the objective, and $\delta_z$ is the axial Gaussian width derived from Eq. (S8-4) and (S8-6). Eq. (S8-7) defines the effective maximum collection angle at each axial depth $z$. Comparison of these theoretical results yields the effective collection angles for the SFN system. By applying this model to the SFN configuration using a 500-μm K9 SMML integrated with a 63×/1.0 NA water-immersion objective under 532 nm excitation, we computed the lateral and axial distributions of the effective signal collection angle (Supplementary Fig. S9).

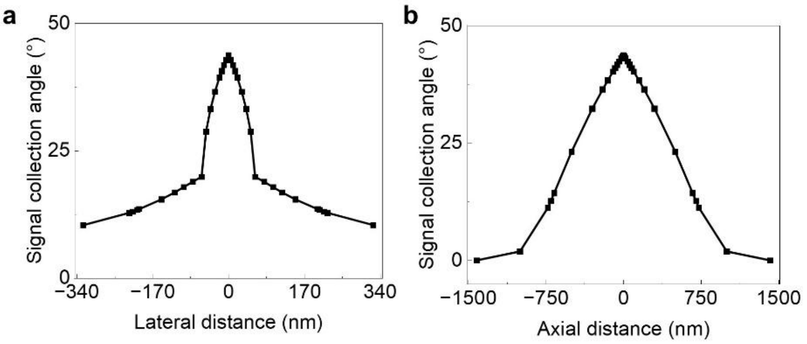

**Supplementary Fig. S9 | Theoretical lateral (a) and axial (b) distributions of signal collection angle of SFN.**

#### 9. Theoretical model of signal-intensity distribution

To compute the lateral and axial signal intensity distributions across the four imaging modalities, WFM, conventional CRM, SIL, and SFN, we decomposed the total signal into excitation and collection components, leveraging the PSF formalism established in Eq. (S8-1).

For the excitation component, the incident laser power is identical in WFM and CRM (both using the same 63×/1.0 NA water-immersion objective and 532 nm excitation); likewise, the effective excitation is identical in SIL and SFN (both employing the same

500-μm K9 SMML). Thus, WFM and CRM serve as the reference excitation baseline, while the excitation PSF for SIL and SFN is scaled relative to CRM by the product of focusing gain $G_f$ and laser-coupling efficiency $\eta_c$. For the specified configuration (500-μm K9 SMML, 1.0 NA objective, 532 nm), this yields:

$$\mathrm{PSF}_{\mathrm{exc,SFN}} = 0.8 \times \mathrm{PSF}_{\mathrm{exc,CRM}} \qquad (S9-1)$$

This scaling reflects the net reduction in peak excitation intensity at the focal plane due to coupling losses ($\eta_c < 1$), partially offset by tighter focusing ($G_f > 1$).

For the signal collection component, the lateral and axial angular distributions of collected signal, $\Omega_{col}$ ($x$,y) and $\Omega_{col}$ ($z$), were derived from the geometric-optics models described in Supplementary Notes 7 and 8, and are presented in Fig. 2k, l. The total detected signal intensity $I_{signal}$ at any spatial coordinate is then given by the product of the local excitation intensity and the corresponding collection efficiency:

$$I_{\mathrm{signal}} \propto \mathrm{PSF}_{\mathrm{exc}} \times \Omega_{\mathrm{col}} \qquad (S9-2)$$

Using Eq. (S9-2), we computed the full two-dimensional lateral (x,y) and one-dimensional axial ($z$) signal intensity distributions for all four configurations. Specifically:

1) Lateral distributions: Normalized excitation PSFs were calculated for each modality (Supplementary Fig. S10a). As expected, WFM and CRM exhibit nearly identical lateral profiles, consistent with their shared objective NA and absence of SMML-induced focusing changes. In contrast, SIL and SFN display narrower lateral excitation distributions due to their higher effective NA ($\mathrm{NA_{eff}}$ = 1.51). Multiplying these

excitation profiles (Supplementary Fig. S10a) by the experimentally calibrated lateral collection angle distributions (Fig. 2k) yielded the final lateral signal intensity distributions (Supplementary Fig. S10b).

2) Axial distributions: Analogously, axial excitation PSFs (Supplementary Fig. S10c) were combined with the axial collection angular distributions (Fig. 2l) to obtain the axial signal intensity profiles (Supplementary Fig. S10d).

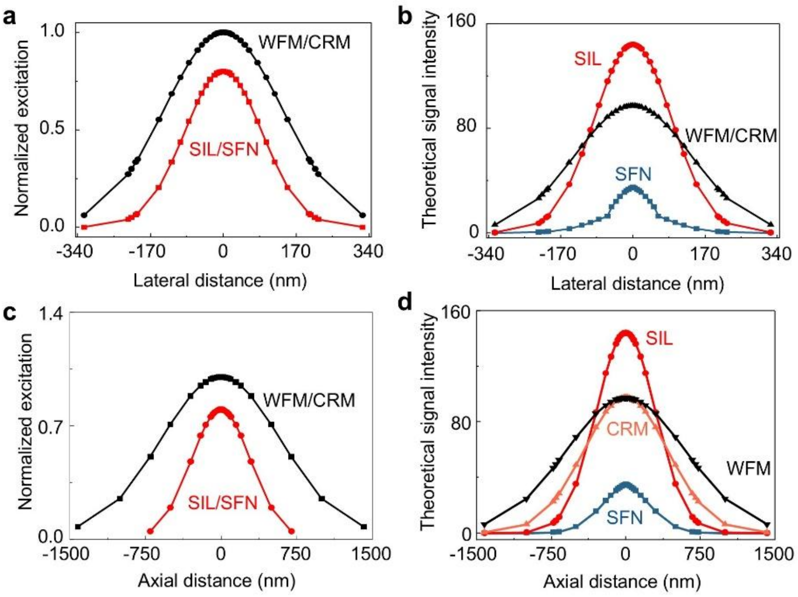

**Supplementary Fig. S10 | Spatial distributions of signal intensity in SFN system. a**, Normalized lateral distributions of excitation intensity for WFM, CRM, SIL and SFN. **b,** Theoretical lateral distributions of Raman signal intensity for WFM, CRM, SIL and SFN. **c**, Normalized axial distributions of excitation intensity for WFM, CRM, SIL and SFN. **d,** Theoretical axial distributions of Raman signal intensity for WFM, CRM, SIL and SFN.

#### 10. Theoretical model for relative signal intensity

To enable quantitative comparison of signal performance across WFM, CRM, SIL, and SFN, we define a normalized metric: the relative signal intensity $S_{total}$. This quantity represents the average power density (integrated signal power per unit area) over the diffraction-limited focal region, specifically, the central Airy disk, and serves as a physically meaningful, system-independent basis for benchmarking signal strength. Mathematically, $S_{\text{total}}$ is defined as the total detected optical power $P_{\text{total}}$ within the focal region, divided by the geometric area $A$ of the central Airy spot:

$$S_{\text{total}} = \frac{P_{\text{total}}}{A} \qquad (\text{S10}-1)$$

For optically opaque, non-scattering, surface-confined samples, which are well approximated as 2D emitters, the lateral intensity distribution $I(x,y)$ is integrated over the central Airy spot:

$$S_{\text{total}} = \frac{\int I(x,y)\, dA}{A} \qquad (\text{S10}-2)$$

Given the rotational symmetry of the Airy pattern about the optical axis, Eq. (S10-2) reduces to a computationally efficient one-dimensional radial integral:

$$S_{\text{total}} = \frac{\int I_x\, dD}{d_{\text{Airy}}} \qquad (\text{S10}-3)$$

where $d_{\text{Airy}}$ is the experimentally measured full-width major-axis diameter of the central Airy spot. For the 3D model, the signal arises from a finite axial extent. In this case, $S_{\text{total}}$ must account for both lateral confinement and axial decay of excitation and collection efficiency. We therefore express the total relative signal intensity as the product of the lateral signal density at the focal plane $S_{\text{lateral}}$ and the normalized axial signal integral:

$$S_{\text{total}} = \frac{1}{2} \times S_{\text{lateral}} \times \int I_z\, dz \qquad (\text{S10}-4)$$

where $\int I_z\, dz$ denotes the normalized axial collection efficiency profile, governed by the combined PSF of excitation and collection, and the integral is taken over the effective axial sampling depth. The factor of 1/2 is a geometric approximation reflecting that only roughly half of the ellipsoidal focal volume lies within the sample when the focus is positioned at the surface. Because this factor is common to all configurations compared, it cancels in relative ratios (e.g., the SBR of Supplementary Note 12) and does not affect the conclusions.

#### 11. Theoretical model of resolution in SFN

According to the PSF, the total lateral resolution of an imaging system is governed by the combined effect of excitation and collection PSFs. When both PSFs are well approximated by Gaussian distributions, their product yields another Gaussian with standard deviation $\delta_{\text{total}}$ satisfying:

$$\frac{1}{\delta_{\text{total}}^2} = \frac{1}{\delta_{\text{exc}}^2} + \frac{1}{\delta_{\text{col}}^2} \qquad (S11-1)$$

Where, $\delta_{\text{exc}}$ and $\delta_{\text{col}}$ are the Gaussian widths of the excitation and collection PSFs, respectively. Converting to FWHM resolution using d = 2.355δ, Eq. (S11-1) becomes:

$$\frac{1}{d_{\text{total}}^2} = \frac{1}{d_{\text{exc}}^2} + \frac{1}{d_{\text{col}}^2} \qquad (S11-2)$$

Where, $d_{\text{total}}$, $d_{\text{exc}}$ and $d_{\text{col}}$ denote the FWHM lateral resolutions of the total system, excitation pathway, and collection pathway, respectively. This relation quantitatively captures how resolution improvement in SFN arises from synergistic enhancement of both excitation confinement (via PRE) and collection angular filtering (via 3D-SFE).

#### 12. Theoretical model of signal-to-background ratio in SFN

The 3D-SFE intrinsic to SFN suppresses out-of-focus and stray light contributions, thereby substantially improving the SBR. To quantify this enhancement, we define the total detected power $P_{\text{total}}$ as the sum of useful signal power $P_{\text{signal}}$ and background power $P_{\text{background}}$:

$$P_{\text{total}} = P_{\text{signal}} + P_{\text{background}} \qquad (\text{S12}-1)$$

Specifically, $P_{\text{signal}}$ is integrated over a defined region of interest (ROI): a lateral disk of radius 100 nm centered on the focal point and an axial slab of thickness 200 nm straddling the focal plane, while $P_{\text{background}}$ represents the background power. The SBR is then defined as:

$$\text{SBR} = \frac{P_{\text{signal}}}{P_{\text{background}}} \qquad (\text{S12}-2)$$

Within the 3D PSF framework, $P_{\text{signal}}$ decomposes into lateral and axial components:

$$P_{\text{signal}} = \frac{1}{2} \times P_{\text{signal,lateral}} \times \int_{-100}^{100} I_z \, dz \qquad (\text{S12}-3)$$

Where, $P_{\text{signal,lateral}}$ is the lateral signal power within the 100-nm-radius disk, and $\int I_z \, dz$ is the normalized axial collection efficiency profile (as defined in Supplementary Note 10). Similarly, the total power integrates over the same axial range:

$$P_{\text{total}} = \frac{1}{2} \times P_{\text{lateral}} \times \int I_z \, dz \qquad (\text{S12}-4)$$

The factor of 1/2 has the same meaning as in Supplementary Note 10 (hemi-spherical bounding of the focal volume by the sample surface) and cancels in the SBR ratio of Eq. (S12-2).

#### 13. Calculation and experimental validation of the effective numerical aperture in SIL and SFN

NA is a fundamental parameter governing both optical resolution and signal intensity in microscopy. For an ideal hemispherical SIL, prior theoretical and experimental studies have established that the $NA_{eff}$ satisfies the relation[2]:

$$\mathrm{NA_{eff}} \approx n_{\mathrm{SIL}} \times \mathrm{NA} \tag{S13-1}$$

Where, $n_{\mathrm{SIL}}$ denotes the absolute refractive index of the SIL material. In the experimental configuration, employing an SMML as the immersion medium, the analogous expression becomes:

$$\mathrm{NA_{eff}} \approx n_{\mathrm{SMML}} \times \mathrm{NA} \tag{S13-2}$$

By combining the conventional definition of NA with the physical interpretation of $NA_{eff}$, we derive the general form:

$$\mathrm{NA_{eff}} \approx n_{\mathrm{eff}} \times \sin \alpha_{\mathrm{eff}} \tag{S13-3}$$

Where, $\alpha_{\mathrm{eff}}$ represents the effective aperture half-angle and $n_{\mathrm{eff}}$ denotes the effective refractive index in SIL system. As predicted by the geometrical-optics model (Supplementary Note 6), $\alpha_{\mathrm{eff}}$ approaches 90° when the SMML is coupled with high-NA objectives (Supplementary Table S1). Under such conditions, Eq. (S13-3) simplifies to:

$$\mathrm{NA_{eff}} \approx n_{\mathrm{eff}} \tag{S13-4}$$

Critically, in both SIL and SFN configurations, the optical path between the objective and the sample comprises two distinct media, the ambient medium (e.g., air or aqueous buffer) and the SMML, not a single homogeneous immersion medium (e.g., oil or water). Consequently, determining $n_{\mathrm{eff}}$ and identifying the parameters that govern its magnitude, is essential for rigorously evaluating and optimizing system performance.

Physically, the enhancement of NA via SIL arises from increasing the effective refractive index of the near-field optical path. For a hemispherical SIL, this enhancement factor approximates $n_{\mathrm{SIL}}$ (cf. Equation S13-2)[2]. In contrast, for a super-hemispherical (Weierstrass-type) SIL, where refraction occurs twice (at both planar and spherical interfaces), the enhancement scales approximately as $n_{\mathrm{SIL}}^2$[3]. In summary, NA of the objective and the refractive index of the SMML jointly determine $\mathrm{NA}_{\mathrm{eff}}$. Accordingly, our analysis focuses on quantifying how these two parameters influence $n_{\mathrm{eff}}$.

We systematically investigated the dependence of $n_{\mathrm{eff}}$ on three variables: NA of the objective, the refractive index of the SMML, and the diameter of the SMML. $\mathrm{NA}_{\mathrm{eff}}$, and thus $n_{\mathrm{eff}}$, was extracted experimentally from the measured FWHM of the Airy spot. In the low-NA regime (0.25–0.55), $n_{\mathrm{eff}}$ increased linearly with objective NA, indicating that the NA of objective imposes the primary limitation. Beyond NA = 0.55, however, $n_{\mathrm{eff}}$ saturated asymptotically toward the bulk refractive index of the SMML (Supplementary Fig. S11 and Supplementary Fig. S12). This saturation behaviour reveals a transition: once the objective NA exceeds a threshold value, the intrinsic refractive index of SMML, not the objective, becomes the dominant limiting factor. Furthermore, $n_{\mathrm{eff}}$ remained invariant across the tested range of SMML diameters

(Supplementary Fig. S13 and Supplementary Fig. S14), confirming that optical performance is governed by refractive index contrast, not physical lens size. Collectively, these results demonstrate that the SIL/SFN system operates in a diffraction-limited regime; upon reaching this limit, $n_{eff}$ is determined almost exclusively by the material properties of SMML.

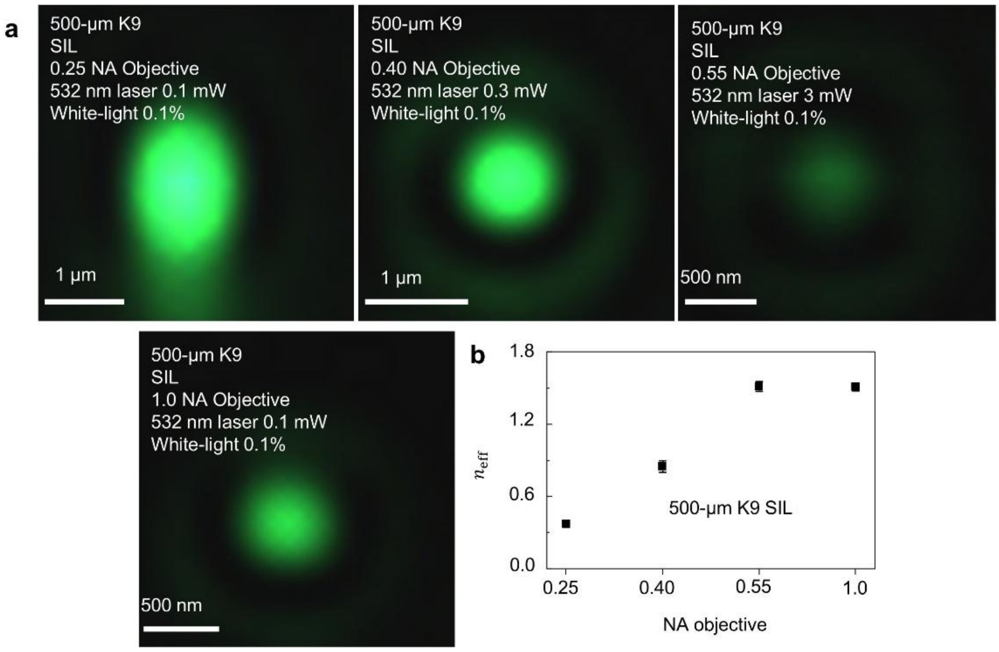

**Supplementary Fig. S11 | Dependence of effective refractive index on objective NA. a**, Images of Airy disk adopting a 500-μm K9 SMML with objectives of different NA (0.25, 0.40, 0.55 and 1.0). **b**, Effective refractive index as a function of objective NA. Data are presented as mean ± s.d. (N = 3).

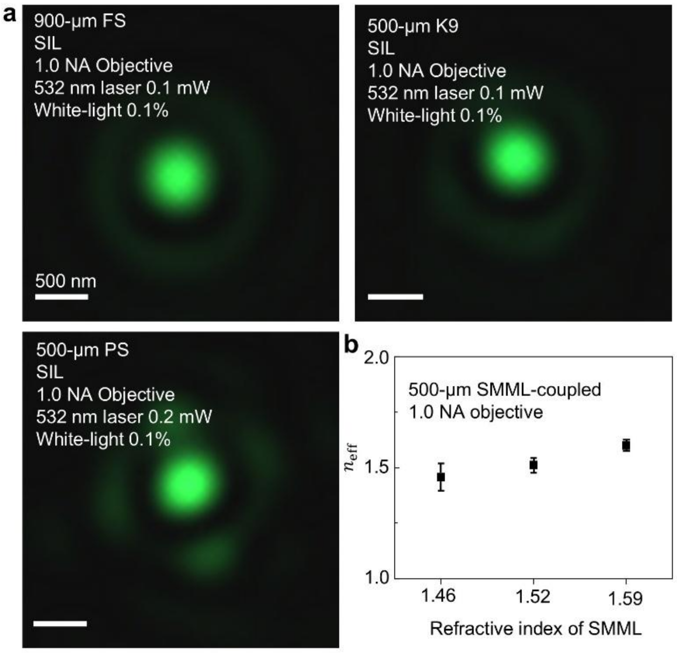

**Supplementary Fig. S12 | Dependence of effective refractive index on the refractive index of SMMLs. a**, Images of Airy disk adopting a 1.0 NA objective and coupled with SMMLs in different refractive indices (FS 1.46, K9 1.52 and PS 1.59). **b**, Effective refractive index as a function of refractive index of SMMLs. Data are presented as mean ± s.d. (N = 3).

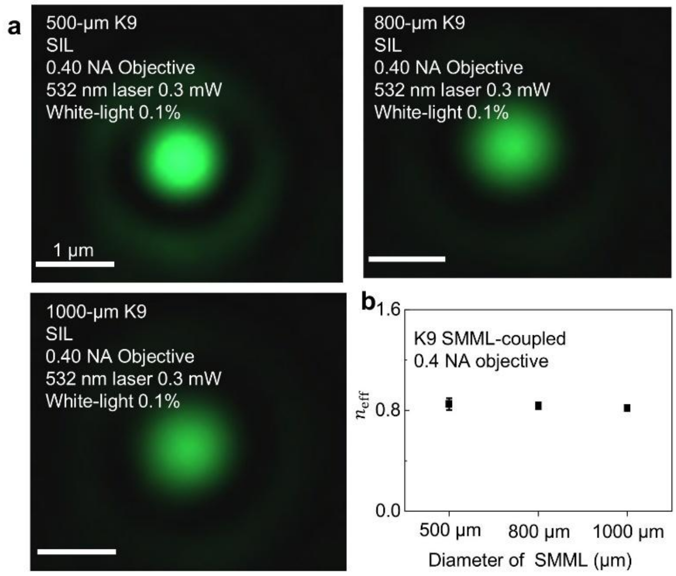

**Supplementary Fig. S13 | Independence of effective refractive index with the diameter of SMMLs. a**, Images of Airy disk acquired via a 0.40 NA objective coupled with K9 SMMLs in different diameters. **b**, Effective refractive index as a function of SMML diameter. K9 SMMLs in diameters of 500, 800 and 1000 μm were selected for the experiments. Data are presented as mean ± s.d. (N = 3).

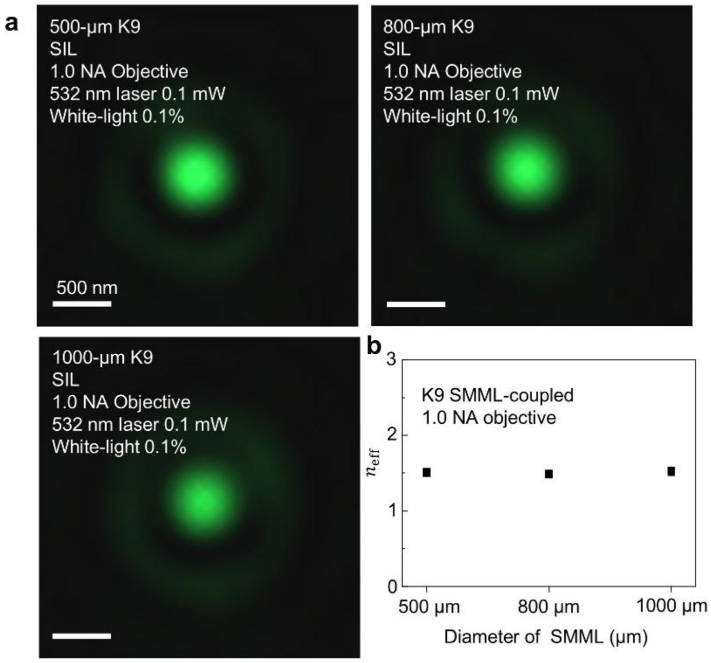

**Supplementary Fig. S14 | Independence of the effective refractive index with SMML diameter. a**, Images of Airy disk acquired via a 1.0 NA objective coupled with K9 SMMLs in different diameters. **b**, Effective refractive index as a function of SMML diameter. K9 SMMLs with diameters of 500, 800 and 1000 μm were selected for the experiments. Data are presented as mean ± s.d. (N = 3).

**Supplementary Table S1| Aperture half-angle of the SIL.**

| Objective | SMML | *R*/μm | *n* | *v*/μm | OD/mm | WD (mm) | Effective OW (mm) | OW (mm) | *α* (°) |
|---|---|---|---|---|---|---|---|---|---|
| 10× | K9 | 150 | 1.52 | 471.0 | 0.321 | 11 | 2.47 | 5.68 | 90 |
| | | 250 | 1.52 | 785.0 | 0.535 | 11 | 2.19 | 5.68 | 90 |
| | | 400 | 1.52 | 1256 | 0.856 | 11 | 1.76 | 5.68 | 90 |
| | | 500 | 1.52 | 1570 | 1.07 | 11 | 1.48 | 5.68 | 90 |
| | FS | 250 | 1.46 | 376.0 | 0.426 | 11 | 0.776 | 5.68 | 90 |
| | PS | 250 | 1.59 | 970.0 | 0.720 | 11 | 3.78 | 5.68 | 90 |
| LD 20× | K9 | 150 | 1.52 | 471.0 | 0.321 | 7.3 | 1.50 | 6.38 | 90 |
| | | 250 | 1.52 | 785.0 | 0.535 | 7.3 | 1.21 | 6.38 | 90 |
| | | 400 | 1.52 | 1256 | 0.856 | 7.3 | 0.787 | 6.38 | 90 |
| | | 500 | 1.52 | 1570 | 1.07 | 7.3 | 0.502 | 6.38 | 90 |
| | FS | 250 | 1.46 | 376.0 | 0.426 | 7.3 | 0.319 | 6.38 | 90 |
| LD 20× | PS | 250 | 1.59 | 970.0 | 0.720 | 7.3 | 2.19 | 6.38 | 90 |
| LD 50× | K9 | 150 | 1.52 | 471.0 | 0.321 | 9.1 | 1.97 | 12.0 | 90 |
| | | 250 | 1.52 | 785.0 | 0.535 | 9.1 | 1.69 | 12.0 | 90 |
| | | 400 | 1.52 | 1256 | 0.856 | 9.1 | 1.26 | 12.0 | 90 |
| | | 500 | 1.52 | 1570 | 1.07 | 9.1 | 0.977 | 12.0 | 90 |
| | FS | 250 | 1.46 | 376.0 | 0.426 | 9.1 | 0.541 | 12.0 | 90 |
| | PS | 250 | 1.59 | 970.0 | 0.720 | 9.1 | 2.96 | 12.0 | 90 |
| 63× | K9 | 150 | 1.14 | 1.33 | 1.14 | 2.1 | 2.83 | 4.77 | 90 |
| | | 250 | 1.14 | 1.33 | 1.14 | 2.1 | 2.90 | 4.77 | 90 |
| | | 400 | 1.14 | 1.33 | 1.14 | 2.1 | 3.00 | 4.77 | 90 |
| | | 500 | 1.14 | 1.33 | 1.14 | 2.1 | 3.06 | 4.77 | 90 |
| | FS | 250 | 1.10 | 1.33 | 1.10 | 2.1 | 3.89 | 4.77 | 90 |
| | PS | 250 | 1.19 | 1.33 | 1.19 | 2.1 | 2.08 | 4.77 | 90 |

LD denotes the long-distance objective; $n$ is the relative refractive index of the SMML; $v$ is the image distance; OW is the objective width; α is the aperture half-angle of the SIL.

#### 14. Material selection for SMML-CRM and Raman peak assignment

To assess the influence of SMML material on spectral fidelity, we compared Raman spectra of single-crystal silicon acquired using SMMLs made of K9, FS and PS (Supplementary Fig. S15a-c). The spectrum obtained with the K9 SMML was nearly identical to that acquired by conventional CRM without an SMML, indicating negligible spectral interference. A similarly small effect was observed for FS. By contrast, PS introduced substantial spectral distortion, demonstrating that the intrinsic Raman response of the SMML material can strongly affect measurement fidelity.

Direct Raman measurements of the SMML materials themselves revealed the origin of these differences (Supplementary Fig. S15d). K9 exhibits a weak Raman feature near 1080 $cm^{-1}$, which is assigned to Si-O stretching vibrations in the silicate glass network. FS shows a broad and weak band mainly in the 500-800 $cm^{-1}$ region, arising from Si-O-Si network vibrations in amorphous silica. In contrast, PS displays strong characteristic Raman peaks across the fingerprint region. The prominent peak near 1001 $cm^{-1}$ is assigned to the benzene ring breathing mode, and the peak near 1602 $cm^{-1}$ corresponds to aromatic C=C stretching vibrations. Additional PS features may also appear in the low-wavenumber and C-H deformation regions. Because these intrinsic PS bands are strong and distributed throughout the fingerprint region, they can overlap with sample Raman peaks, generate false spectral features or obscure weak analyte signals.

These results indicate that K9 is a suitable SMML material for Raman applications because it introduces only weak and non-disruptive spectral features under the present experimental conditions. FS also shows low interference, whereas PS should be used with caution in Raman measurements that require high spectral fidelity.

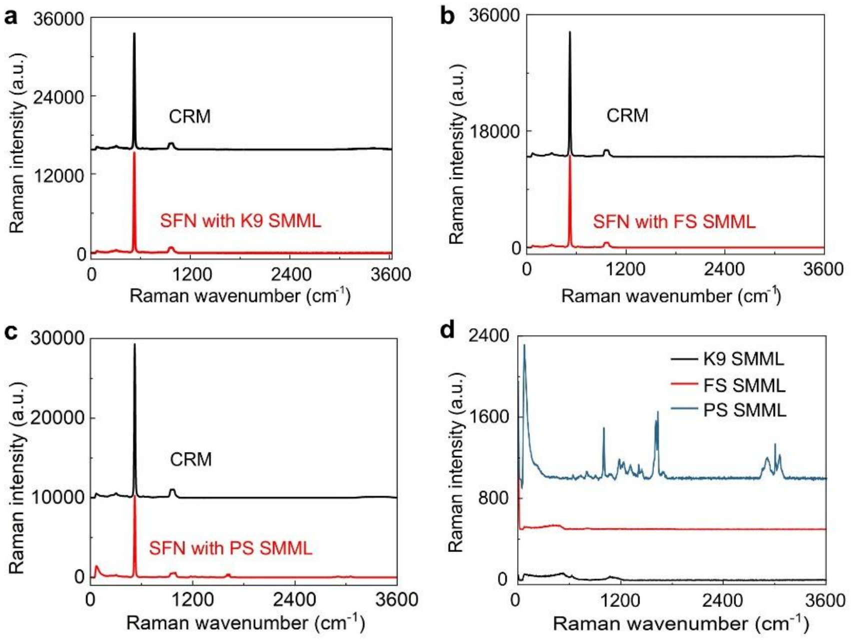

**Supplementary Fig. S15 | Material-dependent spectral interference in SFN. a–c,** Raman spectra of polished Si wafer using SFN with different SMML materials, including K9 (**a**), FS (**b**) and PS (**c**). **d**, Raman spectra of the SMML materials themselves, including K9, FS and PS, showing their intrinsic spectral features. Acquisition parameters: 20 mW laser power, 0.5 s integration time, 10 accumulations.

### Supplementary Figures

#### Supplementary Fig. 1 | Magnification of 500-µm K9 SMML

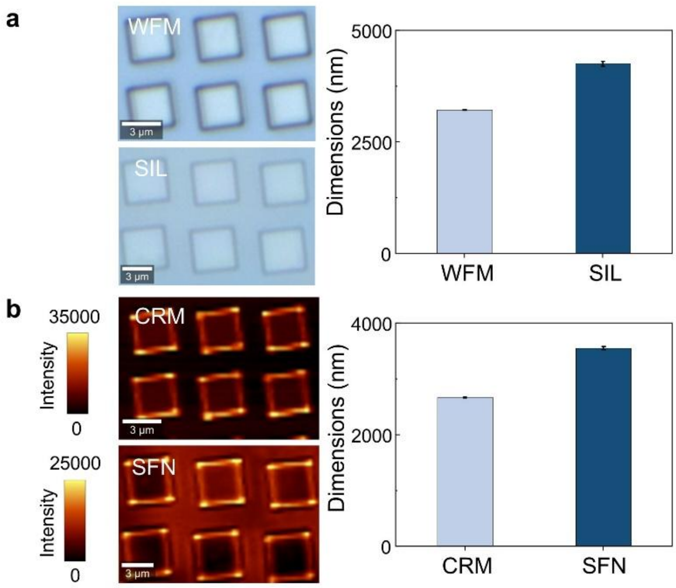

**Supplementary Fig. 1 | Magnification of 500-µm K9 SMML. a**, Bright-field images of the silicon nanostructure obtained using conventional wide-field microscopy (WFM) and solid immersion lens (SIL). **b**, Raman images of the same nanostructure obtained using conventional CRM and SFN. Acquisition parameters of all Raman imaging: 300 nm step size, 520 $cm^{-1}$ Si-Si mode, 0.5 s integration time and 20 mW laser power. All imaging experiments (both with and without the SMML) were performed using the same 63×/1.0 NA objective. Data are presented as mean values ± s.d. over the field of view (FOV).

**Supplementary Fig. 2 | Geometric characterization of the Airy disk in WFM and SIL**

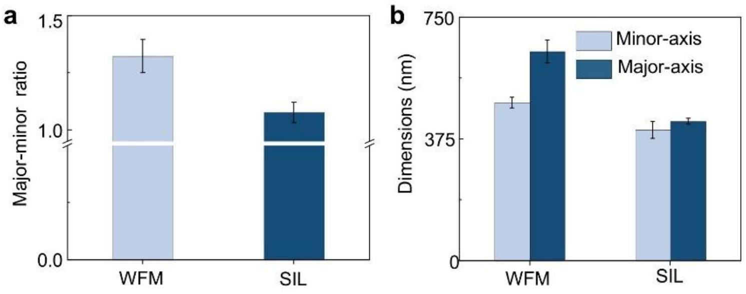

**Supplementary Fig. 2 | Geometric characterization of the Airy disk in WFM and SIL. a**, Ellipticity of Airy disk in WFM system and SIL system. Ellipticity was quantified by the Major-to-minor axis ratio of the Airy disk. **b**, Major- and minor-axis dimensions of the Airy spot in conventional WFM and SIL. Data are presented as mean ± s.d. (N = 5).

**Supplementary Fig. 3 | Analysis of Raman scattering signals in SFN coupled with K9 SMMLs of different diameters**

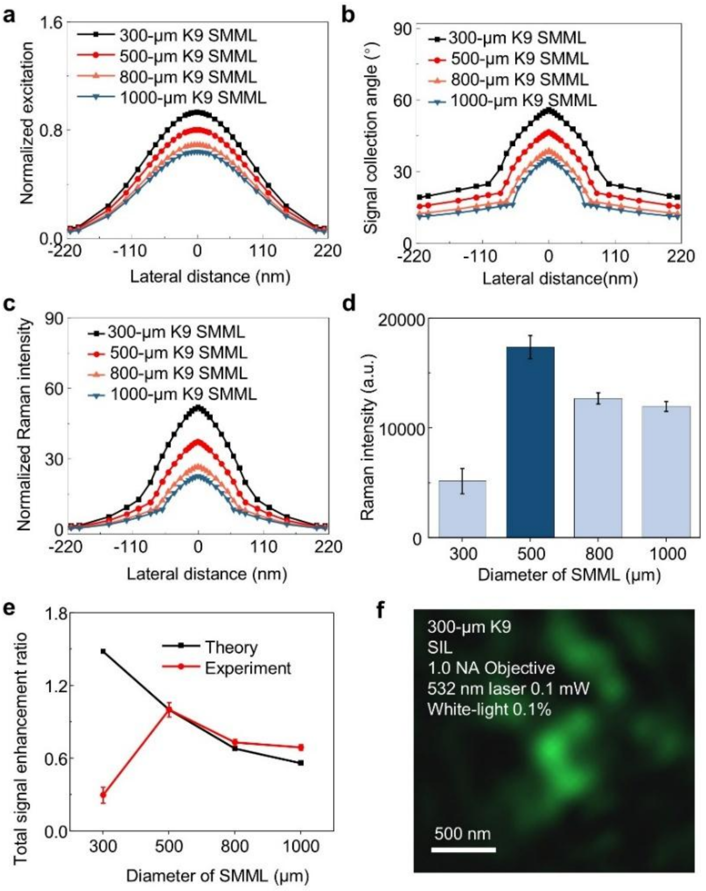

**Supplementary Fig. 3 | Analysis of Raman scattering signals in SFN coupled with K9 SMMLs of different diameters. a-c**, Theoretical lateral distributions in SFN for different diameters of K9 SMML (300, 500, 800 and 1000 μm): normalized excitation intensity (**a**), signal collection angle (**b**) and signal intensity (**c**). **d**, Measured Raman intensity in SFN with different diameters of K9 SMML. **e**, Total theoretical and experimental signal enhancement ratio of SFN with different diameters of K9 SMML. **f**, Airy-disk image acquired under SIL using the 300-μm K9 SMML. Raman signals were measured at the Si-Si stretching mode (520 $cm^{-1}$), under identical acquisition parameters (20 mW laser power, 0.5 s integration time, 10 accumulations). The theoretical values are based on the 2D experimental and theoretical framework established on the polished silicon wafer. Data are presented as mean ± s.d. (N = 5).

**Supplementary Fig. 4 | Analysis of Raman signals in SFN coupled with SMMLs with different refractive indices**

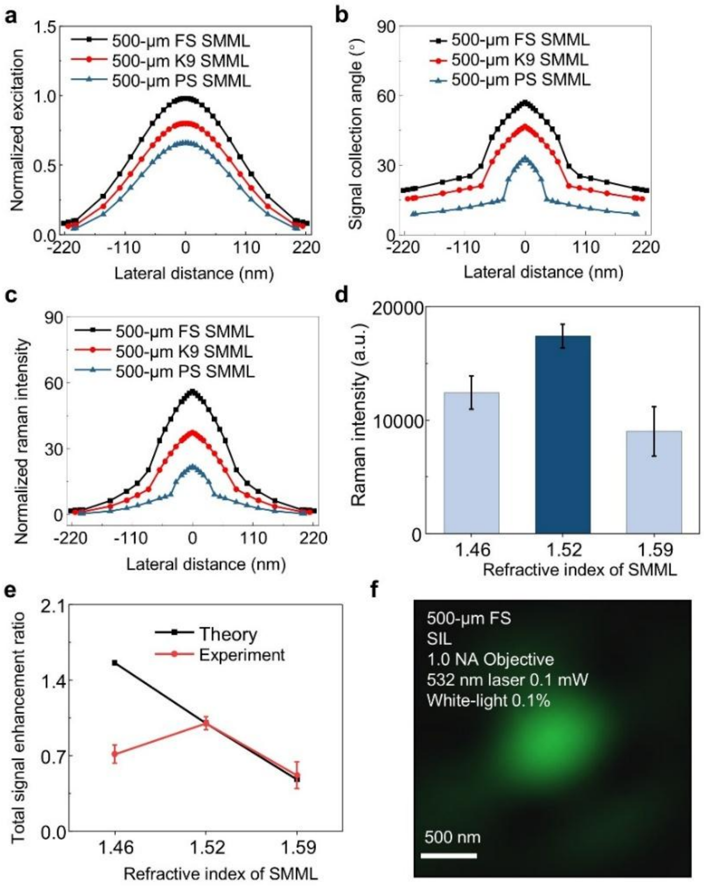

**Supplementary Fig. 4 | Analysis of Raman signals in SFN coupled with SMMLs with different refractive indices. a-c**, Theoretical lateral distributions in SFN for 500-μm SMMLs with different refractive indices (FS, 1.46; K9, 1.52; PS, 1.59): normalized excitation intensity (**a**), Raman signal collection angle (**b**) and Raman signal intensity (**c**). **d**, Raman signal intensity in SFN for different refractive indices of SMML. **e**, Theoretical and experimental total Raman signal enhancement ratio in SFN for different refractive indices of SMML. **f**, Image of Airy disk acquired by SIL using 500-μm FS SMML. Raman signals were measured at the Si-Si stretching mode (520 $cm^{-1}$), under identical acquisition parameters (20 mW laser power, 0.5 s integration time, 10 accumulations). The theoretical values are based on the thin experimental and theoretical framework established on the polished silicon wafer. Data are presented as mean ± s.d. (N = 5).

**Supplementary Fig. 5 | Schematic analogy between an oil-immersion objective and the SMML.**

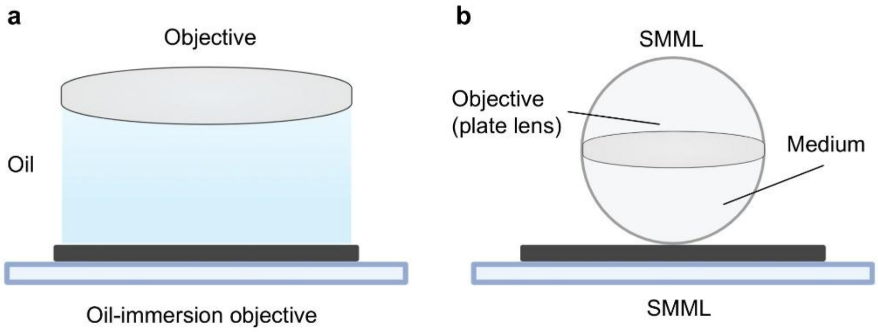

**Supplementary Fig. 5 | Schematic analogy between an oil-immersion objective and the SMML. a**, A conventional oil-immersion objective. The immersion oil acts as a high-refractive-index medium between the specimen and the objective. **b**, An SMML. The SMML serves partly as a high-index immersion medium and partly as an integral optical component of the objective, similar to a plate lens. This schematic illustrates why, under high-NA coupling conditions, the SMML can be regarded as operating in a manner analogous to an oil-immersion objective.

### Supplementary Table

**Supplementary Table 1| Comparison of SFN with representative imaging techniques**

| Technique | labelling | Linearity | Resolution | Key implementation / requirement | Ref. |
|---|---|---|---|---|---|
| STORM | Yes | Yes | ~20 nm | Photoswitchable probes | 4 |
| PALM | Yes | Yes | ~10 nm | Single-molecule localizatio | 5 |
| STED | Yes | No | 35~50 nm | Stimulated-emission depletion | 6 |
| SIM | Yes | Yes | ~100 nm | Structured illumination | 7 |
| ExM | Yes | Yes | ~70 nm | Physical sample expansion | 8 |
| SRS | No | No | Submicrometre | Coherent Raman | 9,10 |
| CARS | No | No | Submicrometre | Coherent Raman | 10,11 |
| S-CARS | No | No | ~1.23× | Saturation-CARS with higher-harmonic demodulation | 12 |
| SSRS | No | No | ~255 nm (1.48×) | Saturation-SRS with harmonic demodulation | 13 |
| Blind-S3 | No | No | ~217 (1.4×) | SRS and blind SIM | 14 |
| HO-CARS | No | No | ~192 nm (1.7×) | Higher-order nonlinear optical signal detection | 15 |
| SREF | Yes | No | ~180 nm | STED and frequency-modulated SREF | 16,17 |
| PSFM-SRS | No | No | ~157 nm (2.2×) | Spatial frequency encoding with π-phase-shift and IFFT | 18 |
| Visible SRS | No | No | ~130 nm | Visible wavelength and high NA objective | 19 |
| URV-SRS | No | No | ~86 nm | Visible SRS, chirping, deep-learning denoising and Fourier reweighting | 20 |
| VISTA | No | No | ~78 nm | Physical sample expansion and SRS | 21 |
| MAGNIFIERS | No | No | ~73 nm | Physical sample expansion and SRS | 22 |
| DO-SRS | Yes | No | ~54 nm | Computational deconvolution | 23 |
| CRM | No | Yes | Submicrometre | Spontaneous Raman scattering | 10,24 |

**Continued Supplementary Table 1| Comparison of SFN with representative imaging techniques**

| Technique | labelling | Linearity | Resolution | Key implementation / requirement | Ref. |
|---|---|---|---|---|---|
| CRM | No | Yes | ~1.4× | Pixel reassignment | [25] |
| SIM | No | Yes | ~1.4× | Structured illumination | [26] |
| SIM | No | Yes | ~1.8× | Structured illumination | [27] |
| Wide-field SIM | No | Yes | ~80 nm | Structured illumination | [28] |
| SFN | No | Yes | ~40 nm | PRE and 3D-SFE | This work |

Techniques are categorized by their physical mechanism and labelling requirements. Abbreviations: SRS, stimulated Raman scattering; CARS, coherent anti-Stokes Raman scattering; SIM, structured illumination microscopy; STED, stimulated emission depletion; STORM, stochastic optical reconstruction microscopy; PALM, photoactivated localization microscopy; ExM, expansion microscopy; S-CARS, saturated CARS; SSRS, saturated SRS; HO-CARS, higher-order CARS; SREF, stimulated Raman excited fluorescence; PSFM-SRS, phase-shifted spatial frequency modulation SRS; URV-SRS, ultrasensitive reweighted visible SRS; VISTA, vibrational imaging of swelled tissues and analysis; MAGNIFIERS, molecule anchorable gel-enabled nanoscale imaging of fluorescence and stimulated Raman scattering microscopy; DO-SRS, deuterium oxide-probed SRS;

**Shading key: Yellow** denotes representative fluorescence-based super-resolution techniques, which rely on exogenous labels and provide high resolution but lack label-free chemical specificity. **Blue** denotes representative nonlinear Raman-based super-resolution techniques, which are label-free but require nonlinear excitation or signal saturation and offer limited resolution gains. **Green** denotes representative linear super-resolution techniques, which operate in the linear regime without nonlinear effects, including linear Raman (CRM) and linear structured illumination approaches; SFN falls into this category as it achieves super-resolution through linear optical signal purification (PRE and 3D-SFE) without nonlinear excitation or signal saturation.
